# Hmox1/CO regulates retina regeneration via modulation of Wnt/β-catenin signalling and induction of *vaspb* in zebrafish

**DOI:** 10.64898/2026.11.02.763727

**Authors:** Rinchen Doma Sherpa, Aditi Sen, Alok Ghosh Chaudhuri, Subhra Prakash Hui

**Affiliations:** S. N. Pradhan Centre for Neurosciences, University of Calcutta, Kolkata 700019, India; Department of Haematology, Nil Ratan Sircar Medical College & Hospital (NRSMC&H), Kolkata -700014, India; Department of Physiology, Vidyasagar College, University of Calcutta, Kolkata 700006, India

**Author notes:** **Corresponding author**: Subhra Prakash Hui, S. N. Pradhan Centre for Neurosciences, University of Calcutta, Kolkata 700019, India., (SPH). **Email addresses of all authors** (RDS), (AS), (AGC).

**Keywords:** zebrafish retina, Müller glia, heme oxygenase, Hmox1/CO signalling, Wnt/β-catenin, Vasp, retina regeneration

## Abstract

Müller glia (MG) in the zebrafish retina retain a latent stem cell capacity that is absent in mammals, yet the injury responsive signals that drive this reprogramming remain incompletely defined. Here, we identify heme oxygenase-1 (Hmox1), the stress-inducible, rate-limiting enzyme of heme catabolism, as a previously unrecognized regulator of this process. Following injury, *hmox1a* is selectively and rapidly induced among the heme oxygenase gene family, localizing to photoreceptors with expression peaking during the MG proliferation window. Pharmacological inhibition and morpholino mediated knockdown of Hmox1 dysregulate the acute inflammatory response, impair Müller glia-derived progenitor cell (MGPC) proliferation, and produce lasting deficits in retinal lamination, establishing Hmox1 as a potentially non-redundant, early-acting node required for structural recovery. Exogenous delivery of carbon monoxide (CO), a bioactive Hmox1 metabolite, partially rescues each of these phenotypes, functionally implicating CO as a key downstream effector. Mechanistically, Hmox1/CO signalling stabilizes β-catenin likely via transcriptional repression of the Wnt antagonist *dkk1b*. This stabilized β-catenin is enriched at a predicted Tcf7l2 site in the promoter of *vaspb*, which encodes an actin binding protein and is newly identified here as an injury-induced Wnt target gene. Bioinformatic and functional analyses further implicate Nfe2l2a (Nrf2) as a candidate intermediary bridging Hmox1/CO to Wnt regulation, with putative antioxidant response elements identified within the *dkk1b* locus. Together, these findings outline a putative Hmox1→CO→Nrf2⊣*dkk1b*→β-catenin→*vaspb* relay coupling redox biology to Wnt-dependent glial reprogramming. Notably, HO-1 is similarly induced in stressed mammalian retina, yet Wnt/β-catenin activation fails to sustain regeneration, positioning the Hmox1/CO axis as a candidate target for future investigation in mammalian retina regeneration.

## Background

Regeneration is a striking exception to the general rule of vertebrate tissue repair, an endogenous restorative program retained by lower vertebrates yet largely lost across mammalian evolution. This capacity relies on an injury-induced microenvironment that reprograms differentiated cells into a progenitor like, multipotent state capable of restoring lost structure and function. Mammals retain a measure of spontaneous regeneration in tissues such as skin and liver (1–3), and rodents and non-human primates mount basal adult neurogenesis following injury (4,5); however, this response falls far short of the structural and functional reintegration achieved in lower vertebrates, and the mammalian central nervous system (CNS), including the retina, remains largely refractory to regeneration.

More than two decades ago, seminal studies by Yurco and Cameron (2005), Fausett and Goldman (2006) and others identified Müller glia (MG) as the resident adult retinal stem cells in zebrafish (6–9). Subsequent single-cell RNA-seq and ATAC-seq analyses of developing and regenerating zebrafish retina, together with cross-species comparisons, have revealed both conserved and species-specific gene regulatory networks governing retinal repair (10–12). These studies converge on a striking principle: although MG enter a comparable reactivated state in both zebrafish and mammals, a distinct regulatory program actively reverts mammalian MG back to a resting state (10), pinpointing the reactivation to reprogramming transition as the decisive, and still poorly resolved, node that separates regenerative competence from regenerative failure.

MG display a conserved radial architecture across vertebrates (13), positioning them as primary sensors of retinal stress and central regulators of tissue homeostasis. During gliosis, MG adopt a pro-inflammatory phenotype, likely triggered by activation of pattern recognition receptors such as Toll-like receptors (TLRs) (14) to shape cytokine output and the earliest phases of neuroinflammation. Notably, the cone-dominant visual processing in zebrafish closely parallels the human retina than rod-dominant rodent models do (15), positioning zebrafish as a physiologically relevant platform for dissecting MG mediated repair with direct translational relevance to human retinal diseases.

Injury elicits a rapid wave of cellular stress across virtually every regenerative paradigm, and this stress response is itself among the earliest triggers of regeneration (16), integrating extrinsic injury cues with intracellular transcriptional programs to switch on cellular reprogramming. Heme oxygenase (HO) is a prototypical stress-responsive gene induced in the CNS under such conditions (17), yet whether it contributes to a regenerative response in the CNS has not been examined.

Heme oxygenase is the rate-limiting enzyme that catabolizes free heme into ferrous iron, biliverdin, and carbon monoxide (CO) (18). Beyond heme turnover, HO-1 is a critical regulator of inflammation and its overexpression protects against inflammatory injury and prolongs xenograft survival (19,20) whereas, HO-1 loss-of-function drives chronic inflammation and immune cell infiltration in both humans and HO-1-null mice (21). HO-2, by contrast, mediates acute inflammatory responses, and its deletion impairs HO-1 induction (22). Heme oxygenase activity has also been implicated in cell proliferation and tumor progression (23,24). Its bioactive metabolite biliverdin functions as a potent antioxidant and anti-nitrosative agent (18), while CO feeds back to upregulate HO-1 (25), alters vascular permeability, and acts as a gaseous neuromodulator in the CNS (26–28).

In the vertebrate eye, expression of both HO-1 and HO-2 is well documented (25,29). HO-2 is constitutively and broadly expressed across retinal neurons (30,31) whereas HO-1 shows stress inducible expression that localizes to Müller glia and photoreceptors in rodents (32,33); pharmacological induction of HO-1 in the eye attenuates inflammation and accelerates wound healing (34,35). In zebrafish embryo, Hmox1 has been previously reported to be expressed in the neuroepithelial cells, a population of polarized developmental stem cells that serve as precursors to the CNS (36) and the heme oxygenase metabolite biliverdin has been shown to regulate NR2E3 during embryonic photoreceptor development (37). However, the expression and function of the heme oxygenase gene family in the injured adult zebrafish retina remain entirely uncharacterized. Given the upregulation of heme oxygenase and its metabolites during stress and injury in the mammalian retina, together with their established roles in inflammation and wound repair, we hypothesized that the heme oxygenase pathway may play a role in orchestrating retina regeneration. Here, we identify a rapid, injury induced upregulation of *hmox1a* in the adult zebrafish retina and propose a previously unrecognized signalling pathway that couples Hmox1/CO output to the core regeneration-associated gene network driving Müller glia reprogramming and retinal regeneration.

## Results

### 1. Zebrafish Heme Oxygenase-1 Is Rapidly and Selectively Induced in the Retina following Stab Injury

To characterize heme oxygenase gene family expression during retina regeneration, we performed a time-course mRNA expression analysis of *hmox1a*, *hmox1b*, *hmox2a*, *hmox2b*, and the biliverdin reductase genes, *blvra* and *blvrb* (Figure 1.A). Among the genes examined, the *hmox1a* paralog was rapidly induced after injury and reached peak expression at 4 dpi in the adult injured retina. Thus, *hmox1a* was selected as the primary candidate for subsequent analysis.

**Figure 1:**
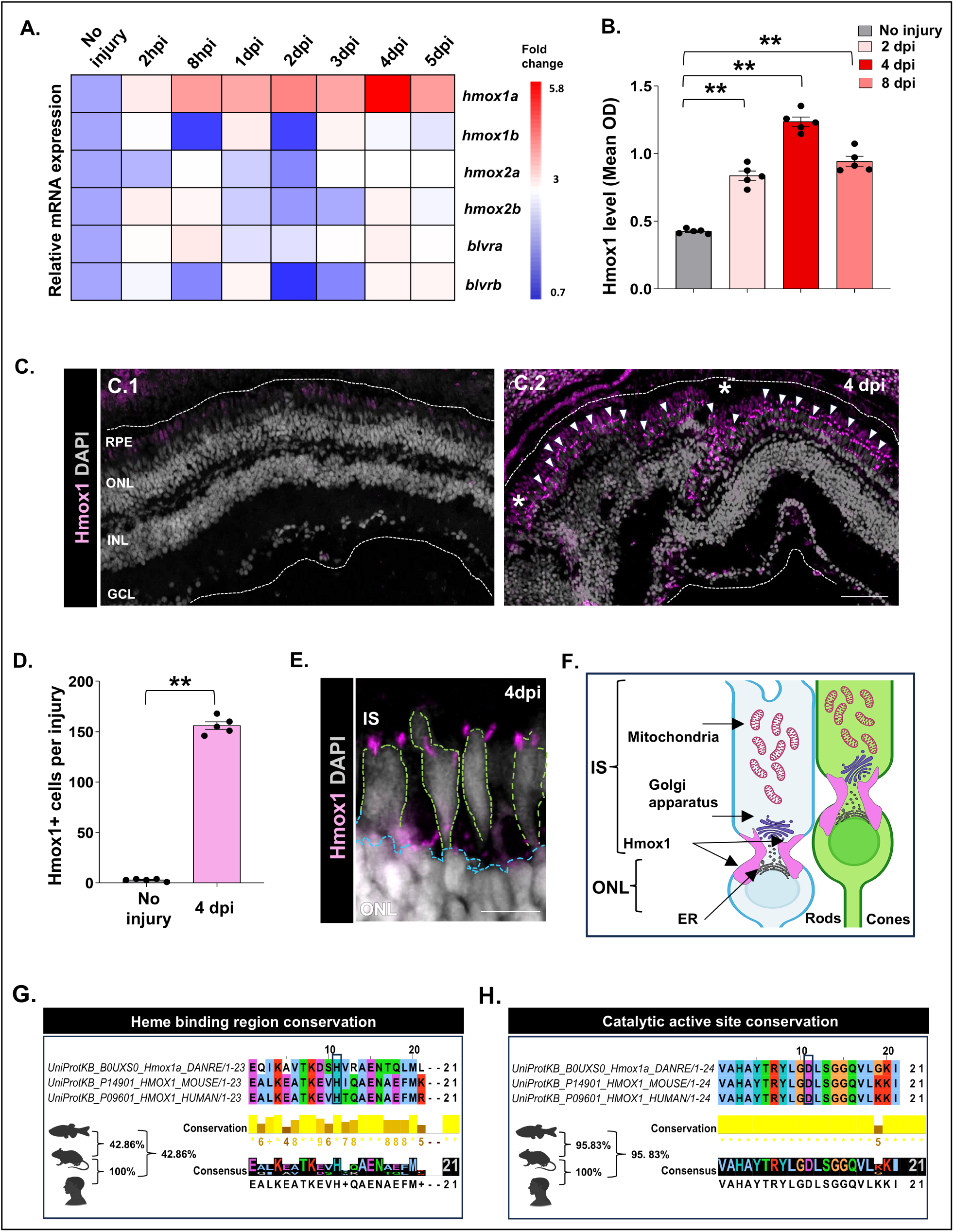
Induction and localization of heme oxygenase expression during zebrafish retina regeneration. (A) qPCR-based heat map showing mean fold change in mRNA expression of *hmox* family (*hmox1a*, *hmox1b*, *hmox2a*, *hmox2b*) and biliverdin reductase genes (*blvra*, *blvrb*) following retina injury in adult zebrafish relative to uninjured controls and normalized to *actb2* (n = 3 biological replicate per time point); (B) Quantitative analysis of Hmox1 protein level at different time points by Indirect ELISA method (Mean ± SEM, n = 5); (C) Immunohistochemical staining of uninjured (C.1) and injured retina (C.2) at 4 days post injury (Transverse Section) showing expression of Hmox1. The white-dotted line extends from the outer nuclear to the ganglion cell layer, white arrowheads mark Hmox1^+^ cells; * represents injury point; Magnification 20X; Scale bar 100 μm; (D) Quantification of Hmox1^+^cells in uninjured and injured retina, (Mean ± SEM, n = 5); (E) Higher magnification of injured retina showing Hmox1 immunoreactivity (pink puncta) on the rod (blue outline) and cone cells (green outline) in the outer nuclear layer at 4 dpi; Magnification 100X, Scale bar 20 μm; (F) Schematic diagram illustrating the proposed cellular and subcellular localisation of Hmox1 within photoreceptors during retina regeneration. Hmox1 is depicted in pink at the nuclear periphery extending toward photoreceptor inner segments; (G-H) Amino acid sequence alignments showing conservation of Hmox1a/HO-1 among zebrafish (*Danio rerio*), mouse (*Mus musculus*), and human (*Homo sapiens*). Rectangles indicate the His25-centered heme-binding region(G) and the Asp140-centered catalytic site (H) respectively; \**p* < 0.05; \*\**p* < 0.01; *ns*, Non-significant; One-way ANOVA with Dunnett’s multiple comparison test (A, B); Mann-Whitney U test (D); RPE= Retinal pigment epithelium, ONL= Outer nuclear layer; INL= Inner nuclear layer; GCL= Ganglion cell layer; IS= Inner segment; ER=Endoplasmic reticulum.

Protein-level analysis by indirect ELISA using an anti-HO-1 antibody confirmed a significant upregulation of Hmox1 in the injured retina at 4 dpi, with a gradual decline at later time points (Figure 1.B). Immunohistochemistry on retinal cryosections (Figure 1.C-D) localized the injury induced expression of Hmox1 to the photoreceptor layer (C.2), with little to no immunoreactivity in the uninjured retina (C.1). High-resolution confocal imaging further revealed a juxtanuclear punctate distribution of Hmox1 in the rod and cone photoreceptors (Figure 1.E-F). Hmox1 immunoreactivity was also detected in the choroid region of the injured eye.

Given the marked injury-induced upregulation of zebrafish Hmox1 during retina regeneration, we next asked whether the key functional motifs in zebrafish Hmox1a are conserved in mammalian HO-1 orthologs. Multiple sequence alignment of two functionally important HO-1 motifs, centred on the heme-binding motif (His25 in human HO-1) and the catalytic active-site motif (Asp140 in human HO-1), revealed 42.86% amino acid identity within the His25-centred motif and 95.83% amino acid identity within the Asp140-centred motif between zebrafish Hmox1a and the corresponding mouse and human HO-1 proteins. Notably, the key functional residues His25 and Asp140 were completely conserved across all three species (Figure 1.G-H).

### 2. Heme Oxygenase-1 Is Required to Restrain the Acute Inflammatory Response and to regulate Müller Glia Proliferation

To investigate the functional role of heme oxygenase 1 during retina regeneration, we pharmacologically inhibited Hmox1 using zinc protoporphyrin IX (ZnPPIX) (Figure 2.A) (36), a heme analogue that competes with heme for the enzyme’s active site thereby blocking Hmox1 and its associated functions. Intravitreal ZnPPIX delivered at 10 μM concentration post-injury significantly suppressed Hmox1 protein levels at 4 dpi (Figure 2.B-D); this effect was corroborated by a corresponding reduction in the number of Hmox1⁺ cells relative to DMSO treated injured controls (Figure 2.B-C). To determine whether inhibition of Hmox1 triggers a transcriptional compensation we profiled *hmox1b* transcripts in ZnPPIX-treated retinas and found no significant induction (Figure S.1), indicating that *hmox1b* is unlikely to compensate for ZnPPIX mediated Hmox1 inhibition in the injured retina (38). Given the well-established role of HO-1 in tempering the magnitude of inflammation (39–41), we next asked whether inhibition of Hmox1 disrupts the early inflammatory window. qPCR profiling of key inflammatory and anti-inflammatory transcripts at 1 dpi and 2 dpi revealed upregulation of *il1b*, *tnfa* and *il6* and a significant downregulation of *il10* and *tgfb* transcripts in ZnPPIX-treated fish, relative to controls (Figure 2. E-F). These findings indicate that Hmox1 inhibition dysregulates the early inflammatory transcriptional response shifting the balance towards a heightened pro-inflammatory signalling and support a role for Hmox1 in modulating the early inflammatory response in the zebrafish retina. In parallel with this hyperinflammatory response, ZnPPIX-treated retinas showed a significant reduction in PCNA⁺ Müller glia-derived progenitor cells (MGPCs) at 4 dpi relative to controls (Figure 2.G-H), linking Hmox1 inhibition to reduced MG proliferation. To analyze long term recovery, multiple injections of ZnPPIX were used, and the retinas were harvested at 28 dpi, a time point at which injured retina achieves phenotypic recovery of retinal lamination (9).

**Figure 2:**
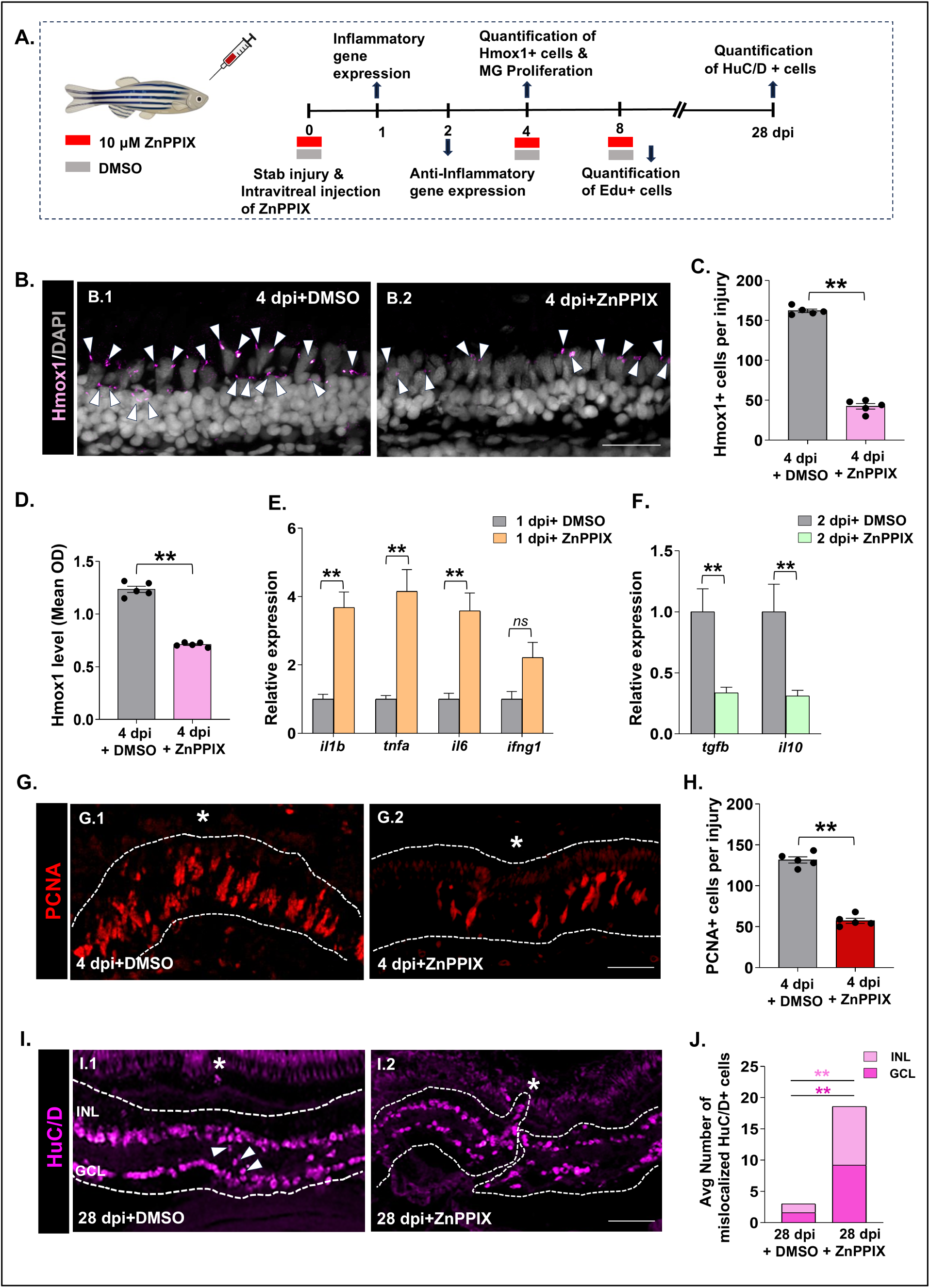
Hmox1 modulates inflammation and is necessary for MG proliferation in injured retina. (A) A schematic timeline for Hmox1 inhibitor treatment showing injury, ZnPPIX treatment and subsequent tissue collections at 1dpi, 2 dpi, 4 dpi, 8 dpi and 28 dpi; (B-C) Immunohistochemical staining and quantification (C) of Hmox1^+^ cells in DMSO (B.1) and ZnPPIX (B.2) treated retina, white arrowheads mark Hmox1^+^ cells, (Mean ± SEM, n=5), Magnification 40X; Scale bar 50 μm; (D) Quantitative analysis of Hmox1 protein level in DMSO and ZnPPIX treated retinas at 4dpi by indirect ELISA method (Mean ± SEM, n=5); (E-F) qPCR-based analysis of inflammatory genes at 1 dpi (E) and anti-inflammatory genes at 2dpi (F) in DMSO and ZnPPIX treated retina (Mean ± SEM, n=5); (G-H) Immunohistochemical staining and quantification (H) of PCNA^+^ cells in DMSO (G.1) and ZnPPIX (G.2) treated retinas at 4 dpi (Mean ± SEM, n=5); * represent injury point; Magnification 40X; Scale bar 50 μm; (I-J) Immunohistochemical staining and quantification (J) of average number of mislocalized HuC/D⁺ cells at the injury epicenter in the INL and GCL in DMSO (I.1) and ZnPPIX (I.2) treated retinas at 28 dpi, (n=5); * represents injury point; white arrow head points towards migrating cells at the injury site; Magnification 40X; Scale bar 50 μm; \**p* < 0.05; \*\**p* < 0.01; *ns*, Non-significant; Mann-Whitney U test; ONL= Outer Nuclear Layer INL= Inner nuclear layer; GCL= Ganglion cell layer.

In striking contrast to DMSO-treated controls, ZnPPIX-treated retinas retained a clearly visible injury site at 28 dpi. HuC/D immunolabeling revealed disorganized lamination with HuC/D^+^ cells displaced from the inner nuclear layer (INL) and the ganglion cell layer (GCL) towards the outer nuclear layer (ONL) and the inner plexiform layer (IPL) (Figure 2.I). Quantification showed a significant increase in the average number of mislocalized HuC/D⁺ cells at the injury epicenter in ZnPPIX-treated fish relative to controls, where mislocalized cells were defined as HuC/D^+^ located at the ONL or IPL (Figure 2.J). Morpholino-mediated knockdown of *hmox1a* downregulated Hmox1 immunoreactivity in the injured retina (Figure S.2) and independently reproduced both the proliferative deficit and impaired long-term recovery (Figure S.3), supporting a requirement for *hmox1a* in these processes.

To distinguish whether the 28 dpi phenotype reflects a sustained requirement for Hmox1 in MGPC proliferation versus a role restricted to the early injury response, we allowed regeneration to proceed unperturbed to 5 dpi before administering ZnPPIX, then assessed EdU incorporation at 8 dpi following a 3-hour pulse. No difference in EdU⁺ cells was observed between DMSO and ZnPPIX treated retinas at this later window (Figure S.4). Together, these data suggest that Hmox1 acts within an early, temporally restricted injury-response phase, and that the impaired structural recovery observed at 28 dpi is consistent with diminished early MG proliferation rather than an ongoing requirement for Hmox1 in progenitor cell-cycle progression.

### 3. CORM-3 Derived Carbon Monoxide Partially Rescues Müller Glia Proliferation and Retinal Recovery following Hmox1 Inhibition

HO-1 and CO participate in a well-documented positive feedback loop; whereby exogenous CO supplementation upregulates HO-1 expression (42–44). In the eye specifically, CO pre-conditioning or post-conditioning elevates heme oxygenase transcripts in porcine retina and confers neuroprotection to retinal ganglion cells against ischemic injury in rat retina (25,45). Building on these precedents, we first tested whether CO could upregulate *hmox1a* in the zebrafish retina. Uninjured retinas were intravitreally injected with 0.9% NaCl, the water-soluble CO-releasing molecule 3 (CORM-3), or its inactivated counterpart iCORM-3 (46), and the transcript levels of the heme oxygenase gene family were assessed one day post-injection. CORM-3 significantly upregulated *hmox1a* at 5 μM, with a non-significant upward trend in the other hmox genes, whereas iCORM-3 produced no significant change in the transcript levels of *hmox* genes unlike CORM-3, indicating a specific, CO-dependent transcriptional response (Figure 3.A).

**Figure 3:**
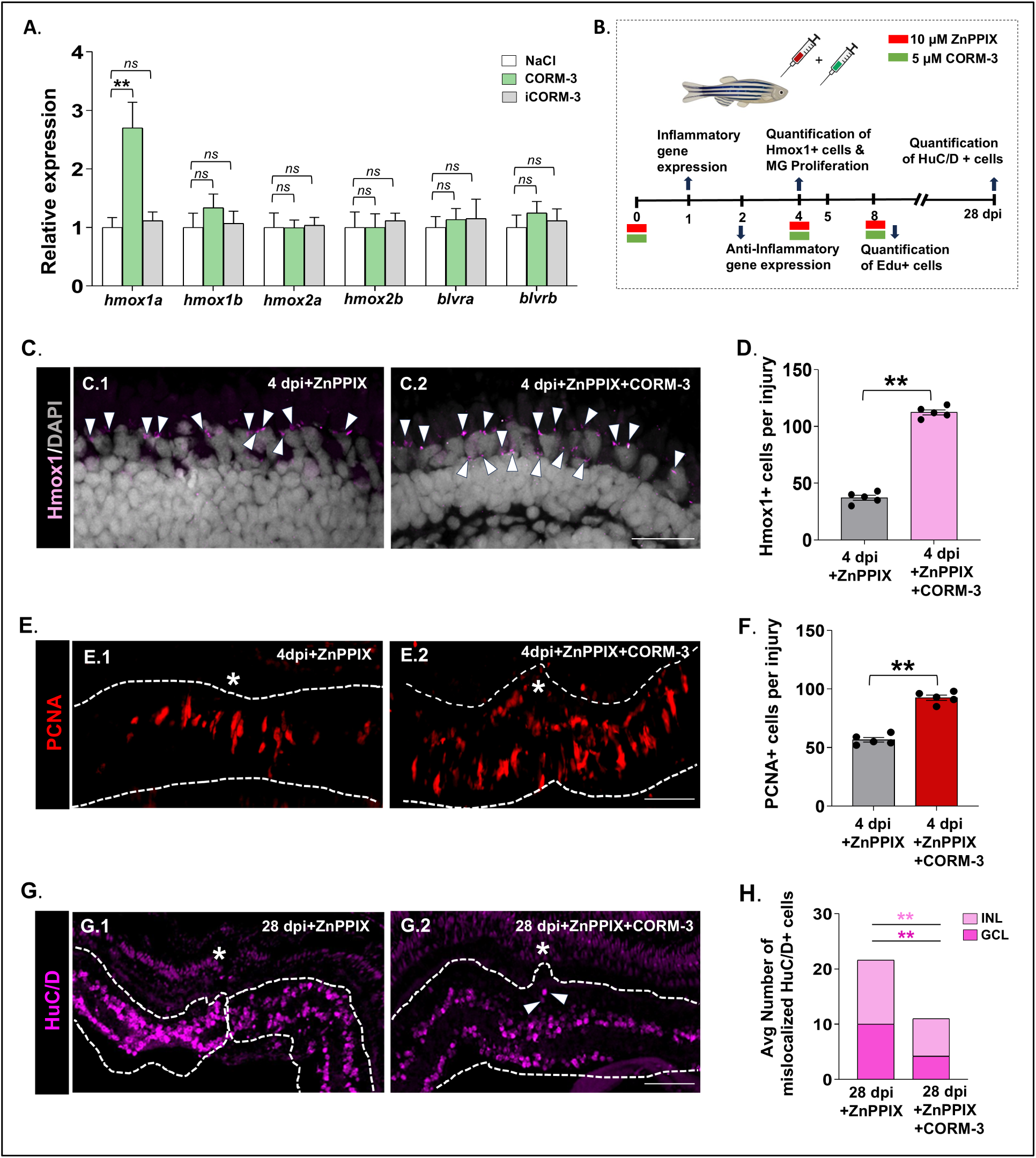
CORM-3 upregulates *hmox1a* transcript levels and partially rescues MG proliferation in the injured retina. (A) qPCR-based analysis of *hmox* family of genes and biliverdin reductase genes (*blvra*, *blvrb*) 1 day after NaCl, CORM-3 and inactivated CORM-3 (iCORM3) treatment in uninjured retina (Mean ± SEM, n = 5); (B) A schematic timeline showing Hmox1 rescue treatment with CORM3 supplementation in ZnPPIX treated retina and tissue collections at 1 dpi, 2 dpi, 4 dpi, 8 dpi and 28 dpi; (C-D) Immunohistochemical staining and quantification (D) of Hmox1^+^ cells in ZnPPIX (C.1) and ZnPPIX +CORM-3 (C.2) treated retina, white arrowheads mark Hmox1^+^ cells, (Mean ± SEM, n=5); Magnification 100X; Scale bar 20 μm; (E-F) Immunohistochemical staining and quantification (F) of PCNA^+^ cells in ZnPPIX (E.1) and ZnPPIX +CORM-3 (E.2) treated retina (Mean ± SEM, n=5); * represents injury point; Magnification 40X; Scale bar 50 μm; (G-H) Immunohistochemical staining and quantification (H) of average number of mislocalized HuC/D⁺ cells at the injury epicenter in the INL and GCL in ZnPPIX (G.1) and ZnPPIX+CORM-3 (G.2) treated retinas at 28 dpi (Mean ± SEM, n=5); * represents injury point; white arrow head points towards migrating cells at the injury site, Magnification 40X; Scale bar 50 μm; \**p* < 0.05; \*\**p* < 0.01; *ns*, Non-significant; Kruskal–Wallis with Dunn’s multiple comparison test (A); Mann-Whitney U test (D, F, H); INL= inner nuclear layer; GCL= ganglion cell layer.

To test whether CO could restore *hmox1a* signalling after ZnPPIX-mediated inhibition, we intravitreally delivered 1 µl CORM-3 one hour after ZnPPIX treatment (Figure 3.B). Supplementation with CORM-3 was able to rescue Hmox1 expression in the injured retina at 4 dpi (Figure 3.C-D). CORM-3 also partially restored MG proliferation at 5 µM, together with the recovery of *hmox1a* transcript levels at this concentration (Figure 3.E-F). To determine whether increasing CORM-3 exposure could further enhance the rescue of MG proliferation, we tested higher concentrations of CORM-3; however, MG proliferation declined at doses above 5 µM (Figure S.5). Given the reported role of CO in regulating cell death (47), and the role of cell death in regulating MG proliferation (48) we assessed apoptosis by TUNEL assay across DMSO, ZnPPIX, and ZnPPIX+CORM-3 treated retinas at 1 dpi and found no significant differences in TUNEL⁺ cell number (Figure S.6), indicating that cell death remained largely consistent across all three conditions and that Hmox1/CO does not regulate MG proliferation through modulation of apoptosis.

Owing to the rescue of MG proliferation, we next asked whether CORM-3 had reversed the hyperinflammatory state induced by Hmox1 inhibition. Exposure to CORM-3 also normalized the expression of *il1b*, *il6* and *tnfa* at 1 dpi relative to ZnPPIX treatment. The anti-inflammatory *il10* and *tgfb* were also significantly upregulated at 2 dpi, indicating that supplementation with exogenous CO can compensate for the loss of Hmox1 activity during the early inflammatory response (Figure S.7). By 28 dpi, CORM-3 exposure also markedly improved recovery of the wounded retinal area, with a decrease in the average number of mislocalized HuC/D⁺ cells at the injury epicenter within the INL and GCL (Figure 3. G-H). Together, these data suggest that CO can upregulate *hmox1a* and partially compensate for Hmox1 inhibition, thereby supporting MG proliferation and structural recovery and implicating the Hmox1-CO signalling in zebrafish retina regeneration.

### 4. Hmox1/CO Signalling Stabilizes β-catenin through Transcriptional Repression of the Wnt Antagonist *dkk1b*

In order to identify the underlying pathways that may act in tandem with Hmox1/CO signalling during retina regeneration, we next examined the levels of regeneration associated genes (RAGs) after Hmox1 inhibition. Candidate genes were selected based on their previously established roles in Müller glia reprogramming, regulation of cell proliferation and retina repair in zebrafish (49–56). Of the 13 RAGs examined, pharmacological inhibition of Hmox1 showed a decreasing trend in the expression of 8 regeneration associated genes (RAGs) (Figure 4.A) with significant downregulation of *vegfa* and *wnt4a*. This expression profile is consistent with the reduced MGPC proliferation observed in our study. Relative to Hmox1 inhibition, CORM-3 treatment showed an increasing trend in RAG expression and significantly restored *wnt4a* levels.

**Figure 4:**
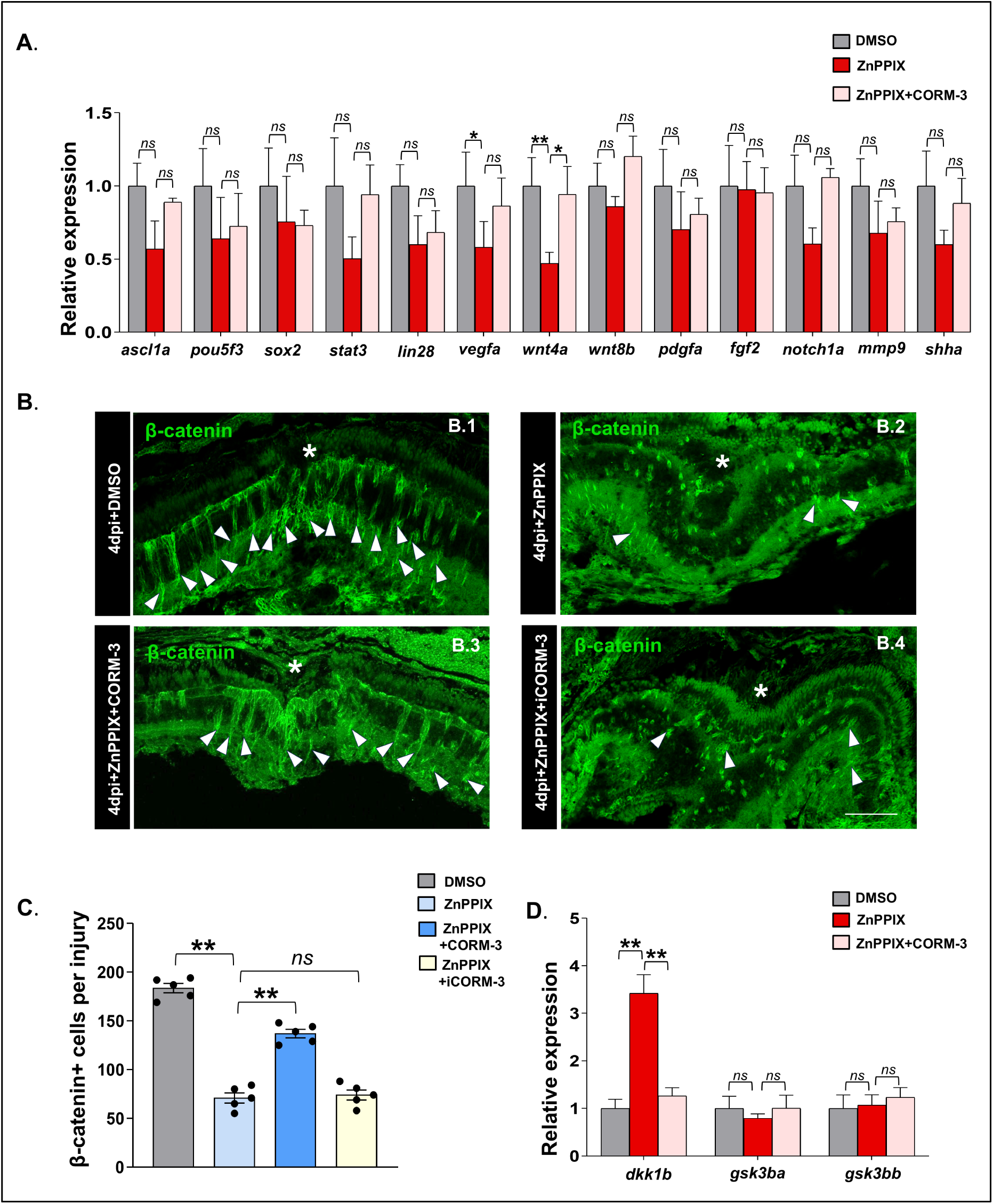
Hmox1/CO modulates β-catenin levels in injured retina. (A) qPCR-based analysis of regeneration associated genes (RAGs) in DMSO, ZnPPIX and ZnPPIX+CORM-3 treated retinas at 4 dpi (Mean ± SEM, n=3); (B-C) Immunohistochemical staining and quantification (C) of β-catenin^+^ cells in DMSO (B.1), ZnPPIX (B.2) and ZnPPIX+CORM-3 (B.3) and ZnPPIX+ iCORM-3 (B.4) treated retinas; * represents injury point; white arrowheads mark β-catenin accumulation at the INL;(Mean ± SEM, n = 5); Magnification 40X; Scale bar 50 μm; (D) qPCR-based analysis of *dkk1b*, *gsk3ba* and *gsk3bb* in DMSO, ZnPPIX and ZnPPIX+CORM-3 treated retina at 4 dpi (Mean ± SEM, n=3); \**p* < 0.05; \*\**p* < 0.01; *ns*, Non-significant; One-way ANOVA with Tukey’s multiple comparison test (A, D), Kruskal–Wallis with Dunn’s multiple comparison test (C).

The Wnt/β-catenin axis is well documented to play a role in injury-induced MG proliferation in both mice and zebrafish (57,58). Given that β-catenin expression is restricted to retinal progenitors in the injured zebrafish retina (57), we examined whether Hmox1 inhibition affected β-catenin expression. Relative to DMSO-treated controls, both ZnPPIX treatment and Hmox1a morpholino knockdown reduced β-catenin expression in the injured retina at 4 dpi (Figure 4.B-C; S.8), paralleling a previous report of reduced active β-catenin in HO-1-knockout mice (59). CORM-3 supplementation in ZnPPIX-treated fish partially restored β-catenin levels, suggesting that the Hmox1/CO signalling also regulates the Wnt/β-catenin signalling in zebrafish retina.

Because Dkk1b and Gsk3-β act as the principal negative regulators of β-catenin stabilization during zebrafish retina regeneration (57), we next profiled *dkk1b*, *gsk3ba*, and *gsk3bb* transcripts across DMSO, ZnPPIX, and ZnPPIX+CORM-3-treated retinas. Inhibition of Hmox1 significantly elevated dkk1b expression whereas, exposure to CORM-3 downregulated *dkk1b* levels. *gsk3ba* and *gsk3bb* transcripts remained unchanged (Figure 4.D). These data suggest that Hmox1/CO signalling stabilizes β-catenin at least in part through transcriptional repression of the extracellular Wnt antagonist *dkk1b*, a negative regulator of β-catenin accumulation during retina regeneration (57).

### 5. Identification of *vaspb* as an Injury-induced Transcriptional Target of β-catenin downstream of the Hmox1/CO Signalling Axis during Zebrafish Retina Regeneration

Vasodilator-stimulated phosphoprotein (VASP) is a member of the Ena/VASP family of actin-binding proteins that localize to sites of active actin assembly and regulate cell migration, adhesion, and filopodia formation (60–62). Earlier investigations have identified *VASP* as a direct downstream target of the Wnt/β-catenin signalling pathway in breast cancer cells (63), and CO has been shown to promote VASP dependent cell migration in human endothelial progenitor cells (64). Given the convergence of Wnt/β-catenin and CO signalling on *VASP*, together with our finding that Hmox1/CO regulates β-catenin levels in the injured retina, we asked whether Vasp functions as a downstream effector of this axis during zebrafish retina regeneration. Of the two *vasp* paralogs, *vaspb* showed a selective injury induced upregulation in the zebrafish retina and was highly upregulated at 4 dpi (Figure 5.A). This was accompanied by a corresponding rise in Vasp protein levels (Figure 5.B). Basal Vasp immunoreactivity in the uninjured retina was largely restricted to the ONL and GCL, whereas at 4 dpi, Vasp expression was upregulated and expanded across the ONL, INL, and GCL layer (Figure 5. C-E). We next checked whether the Hmox1/CO axis regulates Vasp expression in the regenerating retina. Comparison of Vasp protein levels between DMSO, ZnPPIX, ZnPPIX+CORM-3 and ZnPPIX+ iCORM-3 treated retinas at 4 dpi showed that Hmox1 inhibition reduced Vasp expression, while CORM-3 supplementation restored it (Figure 5.F), suggesting Vasp expression is modulated by Hmox1/CO signalling.

**Figure 5.**
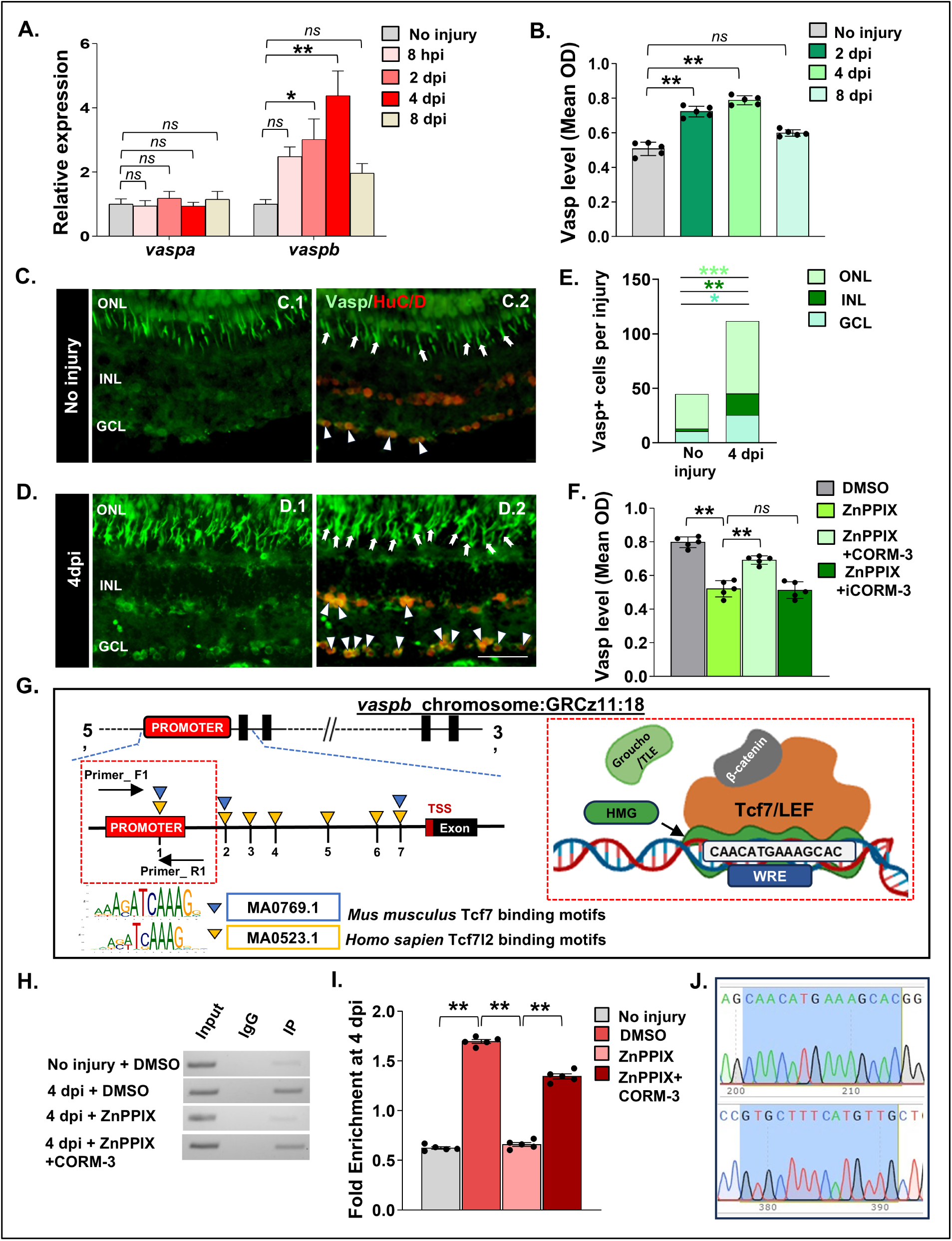
Injury-induced expression and transcriptional regulation of the vasodilator stimulated phosphoprotein (Vasp) during zebrafish retinal regeneration. (A) qRT-PCR analysis of *vaspa* & *vaspb* mRNA, time-course expression in regenerating retina (Mean ± SEM, n=3); (B) Quantitative analysis of Vasp protein level at different time points by indirect ELISA method (Mean ± SEM, n=5); (C-D) Immunohistochemical staining of uninjured retina (C.1-C.2) and injured retina (D.1 –D.2) at 4 dpi (Transverse Section) showing expression of Vasp^+^ cells and co-localized with HuC/D. White arrow-heads point at Vasp^+^ cells in the INL and GCL, white arrows point at Vasp^+^ cells in the ONL, Magnification 40X; Scale bar 50 μm; (E) Quantification of Vasp^+^ cells in the ONL, INL, GCL in injured and uninjured retina at 4dpi (Mean ± SEM, n=5); (F) Quantitative analysis of Vasp protein level at 4 dpi in DMSO treated, ZnPPIX treated, ZnPPIX+CORM-3 and ZnPPIX+iCORM-3 treated retina by indirect ELISA method (Mean ± SEM, n=5); (G) In silico motif analysis identifying Tcf7/Tcf7l2 binding regions within the regulatory regions surrounding the transcription start site (−9 kb upstream to +1 kb downstream) of zebrafish *vaspb* gene (p ≤ 0.0001, q ≤ 0.35). Primers targeting DNA fragments containing the Tcf7/Tcf7l2 motif in the promoter region were used for ChIP–PCR; (H) Chromatin immunoprecipitation (ChIP)-qPCR detection of Tcf7/Tcf7l2 motif on *vaspb*. PCR amplification was carried out with DNA fragments that were immunoprecipitated by anti-β-catenin antibody from injured retina (IP), together with total DNA fragment isolated from uninjured and injured retina (Input) and DNA fragments that were immunoprecipitated by anti-IgG (IgG) antibody from uninjured and injured retina (Mean ± SEM, n=5); (I) ChIP fold enrichment of DNA fragments containing Tcf7/Tcf7l2 motifs was immunoprecipitated using anti-IgG and anti-β-catenin antibody (Mean ± SEM, n=3); (J) Sanger sequencing of ChIP–PCR amplicons confirming identity of the predicted Tcf7/Tcf7l2 binding motif at the *vaspb* promoter region. The conserved motif is highlighted and verified by bidirectional Sanger sequencing (Top -forward, bottom -reverse), \**p* < 0.05, \*\**p* < 0.01; \*\*\**p*< 0.001, *ns,* Non-significant; One-way ANOVA with Dunnett’s multiple comparison test (A, B); Mann-Whitney U test (E), Kruskal–Wallis with Dunn’s multiple comparison test (F); One-way ANOVA with Sidak’s multiple comparison test (I); ONL= outer nuclear layer; OPL=Outer Plexiform Layer; INL= inner nuclear layer; IPL= Inner Plexiform Layer; GCL= ganglion cell layer; TCF7/LEF=T-cell factor/lymphoid enhancer-binding factor; WRE=Wnt-responsive element; TLE= Transducin-Like Enhancer of split; HMG box=High-mobility group box.

Given that Hmox1/CO regulates β-catenin stabilization, and that *VASP* has previously been identified as a Wnt/β-catenin transcriptional target in cancer cells (63), we asked whether β-catenin directly regulates *vasp* transcription in the injured retina. Wnt/β-catenin signalling controls gene expression via the evolutionarily conserved TCF/LEF transcription factor family: nuclear β-catenin complexes with TCF/LEF proteins bound to Wnt-responsive elements (WREs) to activate target gene transcription (65). ChIP-seq studies across multiple human cancer cell lines have identified TCF7L2 (TCF4) as significantly enriched at the *VASP* promoter (63), and ChIP-PCR has confirmed that both β-catenin and TCF7L2 occupy this promoter. Notably, Tcf7l2 is also expressed in the anterior neuroectoderm and optic vesicle during early eye development and has been implicated in eye formation (66). Motivated by these precedents, we searched for Tcf7/TCF7L2 recognition elements within the zebrafish *vaspb* regulatory regions and identified multiple candidate motifs within *vaspb,* including one within the *vaspb* promoter (Figure 5.G; Table S.1).

To test whether β-catenin–Tcf7l2 occupies the *vaspb* promoter during regeneration, and whether this occupancy depends on Hmox1/CO-mediated β-catenin stabilization, we performed ChIP using retinal extracts from uninjured, 4 dpi DMSO, 4 dpi ZnPPIX, and 4 dpi ZnPPIX+CORM-3 treated fish. ChIP-PCR revealed injury-dependent recruitment of β-catenin to the *vaspb* promoter, which was significantly lost upon Hmox1 inhibition and restored by CORM-3 supplementation (Figure 5.H-I); Sanger sequencing of the ChIP-PCR amplicon confirmed the identity of the amplified region containing the predicted Tcf7l2 binding site (Figure 5.J). Collectively, these data identify *vaspb* as a candidate injury inducible transcriptional target of β-catenin and further support a functional interplay between Hmox1/CO signalling and Wnt/β-catenin activity during retina regeneration.

### 6. Bioinformatic and Functional Analyses Identify Nfe2l2a as a Candidate Intermediary Linking Hmox1/CO to Wnt Regulation via Putative AREs within *dkk1b*

To investigate the molecular mechanism underlying Hmox1/CO dependent regulation of β-catenin signalling, we constructed a STRING functional associated network centred on Hmox1a. Using a manually curated gene set based on experimentally validated roles in retina regeneration and documented Hmox1 interactions (67–74), STRING analysis identified predicted interactions between Hmox1a and three members of the Nfe2-like bZIP transcription factor family: Nfe2l2a (Nrf2a), Nfe2l2b (Nrf2b), and Nfe2l3 (Nrf3) (Figure 6.A), consistent with the established role of Hmox1 as a canonical Nrf2 transcriptional target (69). Notably, the network placed Nfe2l2a between Hmox1a and Ctnnb1, nominating Nfe2l2a as a candidate intermediary linking Hmox1a to Wnt/β-catenin signalling (Figure 6.A), consistent with previous reports implicating Nrf2 in Wnt/β-catenin regulation (75,76).

**Figure 6:**
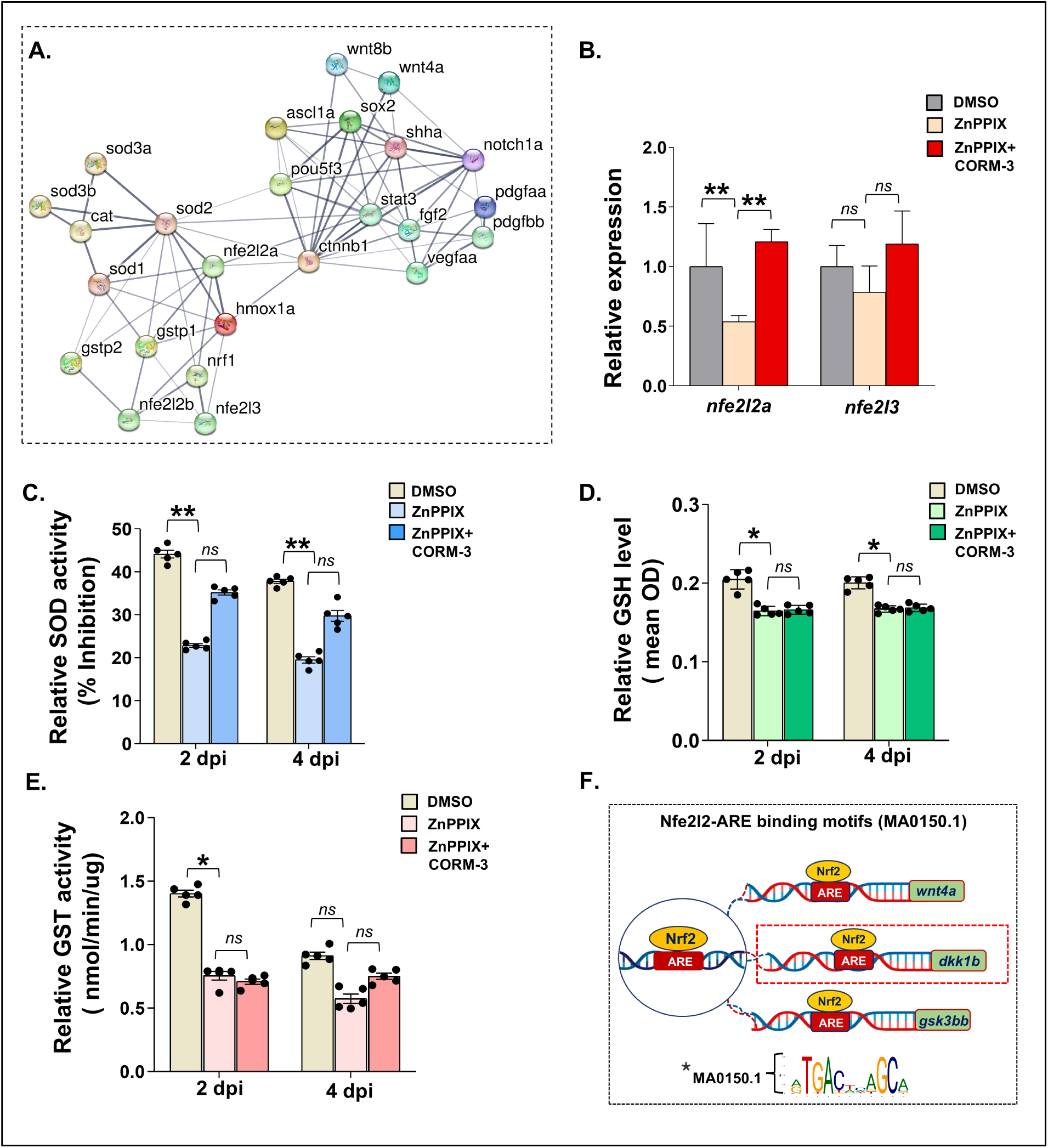
Association of Hmox1/CO and Wnt/β-catenin with NRF2/ ARE mediated antioxidant signalling regulation during zebrafish retinal regeneration. (A) STRING network analysis illustrating predicted protein-protein interactions involving the query protein Hmox1a (red bubble). Nodes are displayed as three-dimensional bubbles representing interacting proteins, while connecting edges indicate confidence based functional associations. Line thickness reflects the strength of supporting evidence (medium confidence threshold = 0.400); (B) qRT-PCR analysis of *nrf2* family in DMSO treated, ZnPPIX treated and ZnPPIX+CORM-3 treated retinas at 4 dpi (Mean ± SEM, n=3); (C) Superoxide dismutase (SOD) activity assessed using pyrogallol auto-oxidation assay at 420 nm and expressed as %inhibition of pyrogallol autoxidation in DMSO treated, ZnPPIX treated, and ZnPPIX + CORM-3 treated retinas at 2 dpi and 4 dpi (Mean ± SEM, n = 5); (D) Reduced glutathione (GSH) levels quantified using DTNB-based spectrophotometric assay at 412 nm. Indirect GSH levels expressed as optical density (OD) in DMSO treated, ZnPPIX treated, and ZnPPIX + CORM-3-treated retinas at 2 dpi and 4 dpi (Mean ± SEM, n = 5); (E) Glutathione S-transferase (GST) activity quantified using a CDNB-based assay with reduced glutathione at 340 nm and expressed as nmol/min/µg protein in DMSO treated, ZnPPIX treated, and ZnPPIX + CORM-3-treated retinas at 2 dpi and 4 dpi (Mean ± SEM, n = 5); (F) Computational identification of Nrf2 (nfe2l2; MA0150.1) responsive antioxidant response elements (AREs) within regulatory regions surrounding the transcription start site (−29 kb upstream to +1 kb downstream) of zebrafish Wnt responsive genes (p ≤ 0.0001, q ≤ 0.25); \**p* < 0.05; \*\**p* < 0.01; *ns,* Non-significant; One-way ANOVA with Tukey’s multiple comparison test (B); Kruskal–Wallis with Dunn’s multiple comparison test (C-E).

Guided by this network, we next profiled the temporal expression of *nfe2l2a*, *nfe2l2b*, and *nfe2l3* in the injured retina. *nfe2l2a* and *nfe2l3*, but not *nfe2l2b*, were significantly upregulated in the injured retina (Figure S.9). Inhibition of Hmox1 significantly reduced *nfe2l2a* in comparison to DMSO treated controls, and rescue with CORM-3 normalized *nfe2l2a* transcript levels (Figure 6.B). In contrast, *nfe2l3* was not significantly affected by Hmox1 inhibition. This finding parallels a prior report in murine cerebral ischemia in which CO-mediated HO-1 induction was Nrf2 dependent and was abolished in Nrf2 knockout mice (76).

Because *nfe2l2a* transcript levels alone do not necessarily reflect Nrf2 pathway activity, we next examined the temporal profile of superoxide dismutase (SOD), reduced glutathione (GSH), and glutathione S-transferase (GST) during regeneration. All three antioxidant readouts peaked at 2 dpi, coinciding with the MG reprogramming window, before declining after 4 dpi (Figure S.10 A-C), and Hmox1 inhibition significantly reduced all three antioxidant readouts (Figure 6.C-E). Of the three, CORM-3 treatment was able to partially restore SOD activity; however, the recovery did not reach significance, whereas GSH and GST remained significantly reduced, suggesting differential CO-sensitivity across antioxidant effectors associated with the Hmox1/CO/Nrf2 axis. The Hmox1/CO dependent partial rescue of SOD is in line with the previous observation that CO exposure increases SOD levels, whereas mice lacking an HO-1 allele were associated with reduction in SOD (77,78). On the other hand, the failure to restore GSH and GST may be attributed to their regulation being dependent on a broader metabolic network (79).

Given that Nrf2 acts transcriptionally by translocating to the nucleus and binding cis-regulatory antioxidant response elements (AREs), we scanned genes with established roles in zebrafish retina regeneration for AREs using the previously identified zebrafish Nrf2-binding motif MA0150.1 (80). Among the identified candidates, *dkk1b*, *wnt4a* and *gsk3bb* contained putative ARE motifs within their regulatory regions (Figure 6.F; Table S.2). Although putative AREs were identified in multiple Wnt-associated genes, the presence of AREs within *dkk1b* was particularly notable, as this finding is consistent with our previous observation that Hmox1 inhibition increased *dkk1b* expression while suppressing β-catenin signalling (Figure 4.D). Collectively, these findings identify Nfe2l2a as a plausible molecular link between Hmox1/CO signalling and β-catenin regulation, with *dkk1b* emerging as a potential downstream transcriptional target during retina regeneration.

## Discussion

While multiple studies have identified key genetic and epigenetic factors that play a critical role in driving retina regeneration, whether the heme oxygenase family of genes contributes to the regenerative response following retina injury has remained largely unexplored.

Among the *hmox* family, our time-course analysis revealed a selective and robust induction of *hmox1a* following stab injury, and the expression was localized to the photoreceptor layer (Figure 1.E). As the primary sensory neurons responsible for converting photons into electrical signals for neurotransmission, photoreceptors are central to retinal function and exhibit exceptional sensitivity to metabolic and oxidative stress (81). Moreover, the retina is highly vascularized in many vertebrates and has one of the highest rates of oxygen consumption per unit weight among the tissues in the body (82). Given these high metabolic and oxidative demands, the injury induced expression of Hmox1 in the photoreceptor layer is therefore consistent with the stress responsive nature of HO-1. In addition, pharmacological inhibition of biliverdin production during zebrafish retina development has been shown to reduce rod and cone photoreceptor populations (37). This induction is also consistent with the prior reports which suggest *hmox1a* to be expressed at substantially higher levels than *hmox1b* during early development and in response to oxidative stress (83,84). At the subcellular level, the juxtanuclear puncta like expression of Hmox1 observed in the photoreceptor layer (Figure 1.E, F) may reflect its association with organelle-rich compartments, such as the endoplasmic reticulum (ER) or mitochondria, both of which are abundant in this region. HO-1 is well established as an ER-resident protein (85) and has also been reported to increase in mitochondria under cellular stress (86). Whether Hmox1 similarly localizes to these compartments in the injured zebrafish retina remains to be determined, and co-localization with ER and mitochondrial markers will be necessary to resolve this question.

Following the identification of *hmox1a* as the predominant injury-responsive paralog, we asked whether its catalytic machinery is conserved. In mammalian HO-1, the His25-centred motif forms part of the heme-binding pocket, where the histidine coordinates the heme iron, and the Asp140-centred motif contributes to the catalytic site that degrades heme (87,88). Amino acid identity across the His25-centred motif was lower than across the Asp140-centred motif (42.86% versus 95.83%; Figure 1.G-H), yet His25 and Asp140 themselves were identical across zebrafish, mouse and human HO-1. Conservation of these two residues suggests that zebrafish Hmox1a retains the core catalytic features of mammalian HO-1.

This injury-responsive induction of *hmox1a* prompted us to investigate its functional significance during retina regeneration. Pharmacological inhibition of Hmox1 dysregulates the expression of *il1b*, *tnfa*, *il6* and decreases *il10*, *tgfb* during the early injury response and is accompanied by reduced MGPC proliferation and impaired long-term restoration of the retinal architecture. These results align with the well established anti-inflammatory function of HO-1 and its catalytic product CO in mammalian tissues (42). Successful regeneration in zebrafish requires an acute but tightly controlled inflammatory response and excessive or prolonged inflammation impairs MG reprogramming and proliferation (89,90). Although *tnfa*, *il1b*, and *il6* each promote Müller glia proliferation (52,91,92), our findings suggest that Hmox1/CO signalling may act as a molecular brake to maintain the balance between inflammatory and anti-inflammatory signalling, which is consistent with the view of the heme-HO system acting as a molecular switch (35) governing the transition from activation to resolution of inflammation during wound healing.

The role of Hmox1 in MGPC proliferation is consistent with previous studies demonstrating a pro-proliferative role for HO-1/CO signalling in both regeneration and cancer. In heterozygous HO-1 knockout rats, hepatocyte proliferation was impaired during liver regeneration, whereas exogenous CO administration promoted regenerative growth after partial hepatectomy via upregulation of hepatocyte growth factor (HGF) (93,94). Similarly in ocular tissues, suppression of HO-1 limits uveal melanoma cell proliferation, whereas HO-1 induction promotes cell-cycle progression (95). However, the reduction in MG proliferation at CORM-3 concentrations higher than 5 μM suggests that the effects of CO on cell proliferation may be concentration dependent, with higher concentrations potentially exerting antiproliferative effects, possibly via modulation of cell death (47,96).

Although a compensatory increase in *hmox1b* expression has been reported in *hmox1a* knockout zebrafish (97), pharmacological inhibition or morpholino knockdown of *hmox1a* did not alter *hmox1b* expression in the injured retina. This difference may arise from distinct cellular responses to stable loss of function mutation when compared to transient disruption of gene function, as deleterious mutations can activate compensatory transcriptional programs that may not be induced by gene knockdown approaches (98). The incomplete regeneration of the retina and the failure to restore HuC/D⁺ neuronal populations at 28 dpi indicate that Hmox1 is necessary for complete structural recovery following injury. The absence of significant changes in the number of EdU⁺ cells at 8 dpi further suggest that Hmox1 may work predominantly during the early response to injury and that the impaired regeneration at 28 dpi may be a downstream effect of an early decrease in MGPC proliferation. Our findings are further supported by a recent study that reports the involvement of Hmox1a in tail fin regeneration in zebrafish larvae (99).

The differential regulation of the Wnt/β-catenin pathway after inhibition of Hmox1 and following CORM-3 rescue points to a functional interaction between Hmox1/CO signalling and canonical Wnt/β-catenin signalling during retina regeneration. Although this interaction has not been investigated previously in the regenerating zebrafish retina, many studies support the existence of a regulatory relationship between HO-1 and canonical Wnt/β-catenin signalling. In a mouse model of postnatal lung repair, genetic deletion of HO-1 reduced active β-catenin levels and impaired tissue recovery following hyperoxic injury, highlighting β-catenin as an important downstream mediator of HO-1-dependent repair (59). The Wnt/β-catenin pathway is well established as a key regulator of injury-induced MG proliferation in both zebrafish and mammals, and in the injured zebrafish retina, β-catenin accumulation in MGPCs serves as a hallmark of retina progenitor identity and active regeneration (53,57,58). The significant reduction in β-catenin immunoreactivity observed in both ZnPPIX-treated and Hmox1a morphant retinas at 4 dpi suggests that the impaired proliferative response following Hmox1 inhibition may be attributed to the dysregulation of the canonical Wnt/β-catenin signalling pathway. The partial restoration of MGPC proliferation and β-catenin immunoreactivity following carbon monoxide exposure provides additional evidence for a role of CO in modulating canonical Wnt/β-catenin signalling during retina regeneration. Interestingly, CORM-3 treatment also restored *hmox1a* transcript levels in our study, suggesting that the partial recovery of β-catenin may be mediated, at least in part, via feedback between Hmox1 and CO. Supporting this idea, a prior study demonstrated that CORM-3 increased HO-1 expression and protected human gingival fibroblasts against inflammatory cytokine-induced loss of β-catenin (100). β-catenin gene transfer has also been shown to activate Wnt signalling and stimulate Müller glia proliferation in the adult mammalian retina even in the absence of retinal injury (58). These observations point to Hmox1/CO as a potential upstream regulator of β-catenin signalling that may modulate MGPC proliferation during retinal regeneration.

The identification of *hmox1a* as the principal functional paralog regulating macrophage recruitment through the MCP-1/CCR2 axis (97), together with reports that macrophages are a major source of Hmox1a at the injury site in zebrafish (97,99), raises the possibility that impaired immune cell recruitment contributes to the incomplete recovery we observed after Hmox1 inhibition. We did not image or quantify macrophages, so this remains untested here. Wnt/β-catenin signalling is itself implicated in monocyte recruitment, with β-catenin transcriptionally regulating CCL2 (MCP-1) (101). Macrophages in turn supply Wnt ligands that promote β-catenin signalling in neighbouring cells (102). A reciprocal relationship at the injury site is therefore plausible, in which Hmox1/CO-dependent β-catenin signalling contributes to macrophage recruitment and recruited macrophages reinforce local Wnt/β-catenin activity. Future studies with macrophage reporter lines combined with Hmox1 inhibition in the retina could help investigate the proposed interaction.

In the regenerating zebrafish retina, Dkk1b has been identified as a negative regulator of canonical Wnt signalling that restricts β-catenin stabilization and MGPC proliferation (57). Indeed, the reciprocal regulation of *dkk1b* following Hmox1 inhibition and CORM-3 supplementation, together with the associated changes in *wnt4a* expression and β-catenin accumulation, supports a model in which Hmox1/CO modulates the Wnt/β-catenin signalling through regulation of *dkk1b* expression.

Although these findings implicate *dkk1b* in Hmox1/CO dependent regulation of canonical Wnt/β-catenin signalling, the molecular mechanism underlying this regulation remains unclear. Our bioinformatic expression analyses identified Nfe2l2a as a potential intermediate linking Hmox1/CO signalling with β-catenin regulation. In response to oxidative stress, Nrf2 works at the transcriptional level by translocating to the nucleus and binding to the cis-regulatory AREs on target genes (80,103). Our bioinformatic analysis identified putative AREs within Wnt-associated genes, including *dkk1b*, *wnt4a* and *gsk3bb*, suggesting that Nfe2l2a may regulate canonical Wnt signalling at multiple levels. However, the reciprocal regulation of *dkk1b* following Hmox1 inhibition and CORM-3 supplementation identifies *dkk1b* as the strongest candidate for Nfe2l2a-ARE regulation. In human cells, a functional ARE in the *DKK1* promoter represses *DKK1* transcription, and mutation of this element abolished the NRF2-dependent regulation (104). The sequence identity of the NRF2 bZIP DNA-binding domain, which mediates ARE recognition, is 73.02% between zebrafish and human (Figure S.10 D). Among the predicted AREs, one was identified near an experimentally annotated zebrafish *dkk1b* transcription start site (TSS), while the other overlapped a predicted promoter region (Figure 7.A). Together these findings support the possibility that an Nrf2-ARE-*dkk1b* regulatory relationship may also be conserved in zebrafish. Suppression of *nfe2l2a* after Hmox1 inhibition, and its rescue by CORM-3, are consistent with the reported ability of CO to release Nrf2 from Keap1 and promote its nuclear translocation and ARE binding (76). Whether Keap1a responds to CO in the same way in the injured retina remains to be determined. These *in silico* predictions require experimental validation, but, combined with the expression data, they support a working model in which the Hmox1/CO-Nfe2l2a pathway regulates canonical Wnt/β-catenin signalling through a transcriptional link to *dkk1b* during retina regeneration (Figure 7.B). Whether Nfe2l2a acts directly on *dkk1b* through these elements remains to be tested by combining *nfe2l2a* knockdown with promoter-reporter assays.

**Figure 7:**
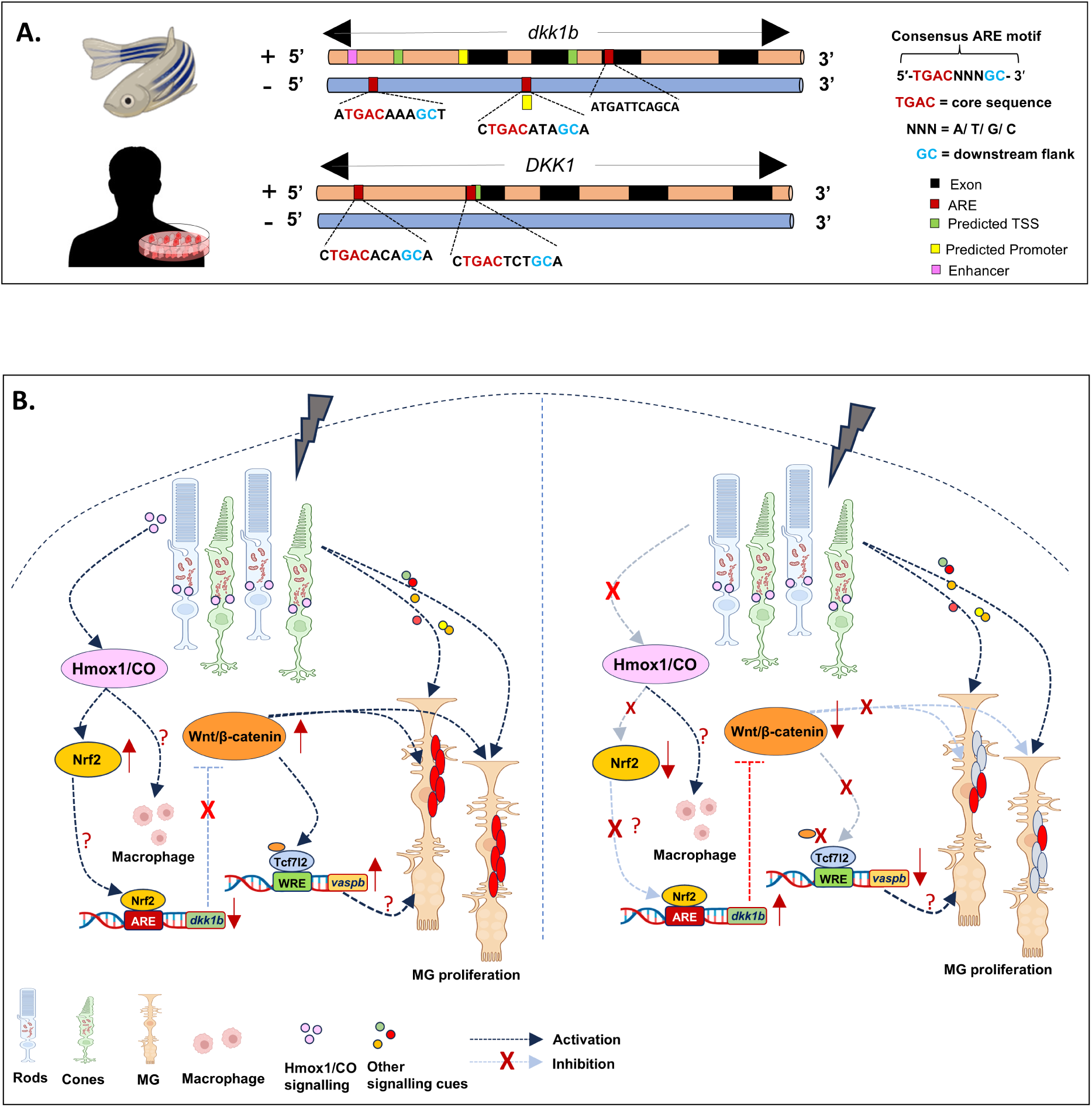
Putative ARE motifs in the *dkk1b* regulatory region and a proposed model for Hmox1/CO signalling during zebrafish retina regeneration. (A) Comparative schematic representation of zebrafish *dkk1b* and human *DKK1* genomic regions. Zebrafish map shows predicted promoter, transcription start site (TSS), and enhancer, with putative ARE motifs positioned relative to the TSS. The consensus ARE motif (5′-TGACNNNGC-3′) is indicated, highlighting conserved regulatory elements across species. (B) A proposed model for Hmox1/CO signalling in zebrafish retina regeneration: Injury activates Hmox1/CO signalling (left), which modulates MG proliferation through the Wnt/β-catenin pathway. Hmox1/CO/Nrf2 may additionally regulate Wnt/β-catenin by modulating *dkk1b*. After injury β-catenin is recruited to the *vaspb* promoter, likely via Tcf7l2, and this is associated with injury-induced *vaspb* upregulation in the zebrafish retina. Inhibition of Hmox1/CO signalling (right) disrupts this signalling cascade and reduces MG proliferation. The roles of macrophages and *vaspb* in this signalling axis remain to be explored.

One of the key findings of the present study is the identification of *vaspb* as an injury-induced gene and a transcriptional target of β-catenin during zebrafish retina regeneration. Vasp immunoreactivity in the injured retina displayed a filamentous staining pattern in the ONL and appeared as ring like structures at the periphery of a subset of cells within the INL and GCL (Figure 5.C-D). Given the established role of Vasp in promoting actin filament assembly, these structures could potentially represent sites of dynamic actin organization (105,106), although this will require co-localization with F-actin markers. The identity of Vasp^+^ cells and whether their localization is associated with cell migration during retina regeneration remain to be explored.

β-catenin-dependent regulation of VASP was first reported in human breast cancer cells (63); our results extend this regulation to an *in vivo* model of tissue regeneration. Wnt/β-catenin signalling controls transcription through the evolutionarily conserved TCF/LEF family (107). Our ChIP-PCR experiments revealed injury-dependent enrichment of β-catenin at the predicted Tcf7l2-binding region within the *vaspb* promoter. Importantly, Sanger sequencing of the ChIP-PCR amplicon obtained following immunoprecipitation with the β-catenin antibody confirmed the identity of the predicted Tcf7l2-binding region within the *vaspb* promoter. Moreover, β-catenin enrichment at this region was reduced following Hmox1 inhibition and restored by CORM-3 supplementation, indicating that β-catenin occupancy at the *vaspb* promoter is dynamically associated with Hmox1/CO signalling. TCF/LEF proteins carry a conserved N-terminal β-catenin-binding domain and switch between repression and activation according to Wnt input (65). An additional effect of Hmox1/CO on Tcf7l2 therefore cannot be excluded. Together, these findings support a model in which Hmox1/CO signalling promotes β-catenin occupancy at the *vaspb* promoter and contributes to the injury-induced transcription of *vaspb* during zebrafish retina regeneration. Interestingly, HO-1/CO has also been shown to regulate VASP post-translationally in mouse endothelial cells. Deletion of HO-1 reduced VASP phosphorylation and exogenous CO induced phosphorylation in both HO-1^+/+^ and HO-1^-/-^cells, consistent with HO-1-derived CO working upstream of VASP phosphorylation (108).

To our knowledge Vasp phosphorylation has not yet been investigated in zebrafish. Given the previous evidence that HO-1-derived CO promotes VASP phosphorylation, Hmox1/CO signalling may regulate Vasp phosphorylation in addition to the β-catenin-dependent transcriptional regulation of *vaspb* observed in our study. Whether Hmox1/CO pathway contributes to Vaspb phosphorylation in the regenerating zebrafish and whether this phosphorylation is functionally linked to actin cytoskeletal remodelling or progenitor migration requires further investigation.

## Conclusion

Collectively, these findings broaden the current understanding of the molecular network underlying zebrafish retina regeneration by identifying injury-induced expression of *hmox1a* and demonstrating a role of Hmox1/CO in the regulation of canonical Wnt/β-catenin signalling. Within this signalling axis, we further implicate Nfe2l2a to be a potential regulator of *dkk1b* (Figure 7.B). Notably, HO-1 is similarly induced under stress and in neurodegenerative conditions (32,109) in the mammalian retina, however the resulting activation of Wnt/β-catenin post injury is transient and does not sustain MG proliferation (58). Taken together with our findings in zebrafish, this raises the possibility that dysregulated Hmox1/CO/β-catenin signalling may contribute to the failure of mammalian Müller glia to sustain a regenerative state, positioning the Hmox1/CO axis as a candidate target for future investigations in the mammalian retina. Moreover, the identification of *vaspb* as a direct transcriptional target of β-catenin identifies a previously unrecognized regulatory connection between canonical Wnt/β-catenin signalling and *vaspb* during retinal regeneration.

## Materials and Methods

### Experimental model, retina injury and pharmacological reagents

All experiments were performed on adult zebrafish (AB strain) between ages of 4-6 months maintained at 28°C under a 14 h light/10h-dark cycle. Zebrafish were anesthetized with tricaine methane sulfonate and subjected to retinal stab injury (7) with minor modifications. Briefly, the right eye was carefully pushed out from its socket and a sterile 31G needle was used to puncture the sclera four times, once in each quadrant. The needle was inserted up to the bevel’s length (∼0.5 mm) for each poke. All experimental procedures were conducted in compliance with the animal ethics guidelines approved by the University Committee for the Purpose of Control and Supervision of Experiments on Animals (CPCSEA).

ZnPPIX, CORM-3 and iCORM-3 were intravitreally injected in the zebrafish eye using a Hamilton syringe. 1 mM stock solution of ZnPPIX (Caymanchem) was made in DMSO, CORM-3 (Sigma Aldrich) was dissolved in 0.9% NaCl and freshly prepared at 0.5 mM stock concentration before use (Key Resource Table). iCORM-3 was generated by incubating CORM-3 at room temperature for 48 hr for spontaneous CO release. For all short-term experiments a single injection of ZnPPIX followed by CORM-3 was used however, for long term phenotype recovery three injections were administered at 0 dpi, 4 dpi and 8 dpi.

### Morpholino application

*hmox1a* antisense Vivo-Morpholino (5′-TGGAGTCCATCTTTGTGCTGTAGA -3′) and a standard control Vivo-Morpholino (Gene Tools LLC) were used for gene knockdown. Vivo-Morpholino delivery to the retina was performed as previously described (110). Approximately 1.0 μl of vitreous humor was withdrawn from the eye using a Hamilton syringe prior to injection, allowing more direct contact between the injected solution and the retina. A 40 μM working solution of Vivo-Morpholino was then intravitreally injected into the zebrafish eye.

### qRT-PCR

Total RNA was extracted from zebrafish retinas using TRIzol reagent (Ambion) and the GoScript™ Reverse Transcriptase Kit (Promega) was used for cDNA synthesis. Gene specific primers have been listed in the oligonucleotide table. qRT-PCR was performed in triplicates with SYBRTM Power Master Mix (Applied Bio systems, USA) and the mRNA levels were normalized to β-actin2.

### Immunohistochemistry

Eyes were enucleated and fixed in 4% paraformaldehyde at 4°C overnight. Injured and uninjured eyes were embedded in O.C.T. compound and cryo-sectioned at a thickness of 10 μm. For immunohistochemistry, slides containing retinal cryosections were rehydrated and washed with PBST, followed by antigen retrieval in sodium citrate or Tris buffer at 80 °C for 5–7 min. Sections were then blocked for 1 h with 5% horse serum and 1% BSA. Sections were incubated with primary antibody for 1 h at room temperature or overnight at 4°C, followed by PBST washes and incubation with secondary antibody for 2 h at room temperature. Nuclei were counterstained with DAPI, and sections were mounted with Gelvatol and stored in the dark. All antibodies are listed in the Key Resource table. According to the supplier’s sequence analysis the anti-HO1 antibody is predicted to recognise Hmox1a with no significant similarity of the immunogen to Hmox1b.

### EdU Labelling & TUNEL assay

The proliferating cells were labelled using the Click-iT™ EdU Cell Proliferation Kit for Imaging, Alexa Fluor™ 488 dye (Invitrogen) at 8 dpi. EdU labelling was performed by intraperitoneal injection of a mixture of 10mM EdU and 3.5% sodium bicarbonate made in tank water 3 hours before collection at 8 dpi.

TUNEL (terminal deoxynucleotidyl transferase dUTP nick end labelling) was performed at 1 dpi on retinal cryosections using the *In situ* Cell Death detection Kit, Fluorescein (Roche) according to the manufacturer’s instructions.

### Microscopy

IHC on retina sections were photographed with an Olympus BX53F2 microscope. Confocal images were captured with an Olympus FLUOVIEW FV3000 confocal laser scanning microscope. Images were processed using FV31S-SW software (Olympus) and Fiji/ImageJ (NIH). The cells were quantified with Cell counter plugin (ImageJ).

### ELISA

Zebrafish were dark adapted overnight and the retina was isolated as previously described (111). Retina protein lysates were prepared by pooling four retina per sample and five independent pools were collected for each experimental condition (n=5) and 20 µg of total protein was loaded per well. Retina were homogenized in RIPA buffer and protein extracts were coated on 96-well plates overnight at 4°C. Wells were blocked and then incubated with primary and HRP-conjugated secondary antibodies and developed with TMB substrate (112). The reaction was stopped with 1 M H_2_SO_4_ and the absorbance was taken at 450 nm using Varioskan LUX multimode microplate reader (Thermo Fisher scientific).

### Antioxidant activity assay

For SOD, GSH and GST activity assay retina were pooled (n=10), dissected and homogenized in 0.1M phosphate buffer (pH 7.4) containing 1Mm EDTA and centrifuged at 13000 rpm for 20 minutes at 4^0^C. SOD activity was measured by the pyrogallol autoxidation inhibition method (113). The absorbance was recorded at 420 nm at 30 sec intervals for 5 min and SOD activity was calculated as percent inhibition of pyrogallol relative to controls. GST activity was determined using the substrate 1-chloro-2, 4-dinitrobenzene (CDNB) and reduced GSH as co-substrate (114). The absorbance was monitored at regular intervals at 340 nm for 5 minutes. The GSH levels were determined by Ellman’s method (115) followed by deproteinized by 5% trichloroacetic acid (TCA) and the supernatant containing GSH was reacted with 5,5’-Dithiobis-(2-nitrobenzoic acid) (DTNB). Absorbance was measured at 412 nm. The protein concentration of retinal tissues lysates was determined by Bradford assay and the volumes of sample per well were normalized to 20ug protein per well.

### Bioinformatic Analysis

STRING was used to analyze protein-protein interaction networks. The transcription factor binding motif sequence of TCF7L2 was downloaded from JASPAR. Upstream regulatory sequences of candidate genes were retrieved from Ensembl and scanned for putative transcription factor binding sites using FIMO-MEME Suite. Protein sequences were retrieved from UniProt and aligned with Clustal Omega (EMBL-EBI), and sequence alignments were visualized using Jalview.

### Chromatin Immunoprecipitation & ChIP-PCR

Chromatin was isolated from dark adapted adult zebrafish retina using EpiQuik™ Tissue Chromatin Immunoprecipitation (ChIP) Kit (Epigentek, P-2003-2). For each condition chromatin was prepared from pooled sample of 14 retina (n=3). Tissues were isolated from control and treated fishes at 4 dpi and fixed in 1% Paraformaldehyde and washed with 1.25M glycine, the tissues were then homogenized using a Dounce homogenizer and centrifuged to remove supernatant. The homogenized tissue was lysed in a buffer containing protease inhibitors, and the chromatin was sheared by sonication to obtain DNA fragments of approximately 200– 1000 bp. The chromatin was immunoprecipitated with anti-β-catenin antibody and non-immune IgG was used as negative control. The immunoprecipitated complexes were washed; cross links were reversed at 65°C and DNA was purified using spin columns. The elute was used for ChIP-PCR amplification of *Tcf7l2* binding region on the *vaspb* promoter and the presence of *Tcf7l2* binding motif was confirmed with Sanger sequencing.

### Statistical Analysis

Values are expressed as mean ± S.E.M. Statistical significance was determined using the Mann–Whitney U test or Student’s t test for pairwise comparisons. For multiple groups and conditions One-way ANOVA with Dunnett’s multiple comparison test and the Kruskal–Walli’s test with Dunn’s multiple-comparisons test were used. Statistical significance was taken at \**p* < 0.05; \*\**p* < 0.01. qRT-PCR analysis was performed on 3-5 biological replicates.

## Abbreviation

MG: Müller Glia
scRNA seq: single cell RNA sequencing
ATAC seq: Assay for Transposase-Accessible Chromatin
TLRs: Toll-like receptors
HO: Heme oxygenase
CO: Carbon monoxide
ZnPPIX: Zinc Protoporphyrin IX
MGPCs: Müller Glia progenitor cells
il1b: Interleukin 1 beta
il6: Interleukin 6
ifng1.2: Interferon gamma
tnfa: Tumor necrosis factor alpha
il10: Interleukin 10
tgfb: Transforming growth factor beta
DMSO: Dimethyl sulfoxide
ONL: Outer Nuclear Layer
INL: Inner Nuclear Layer
GCL: Ganglion Cell Layer
EdU: 5-ethynyl-2’-deoxyuridine
CORM-3: Carbon monoxide releasing molecule 3
iCORM-3: Inactivated carbon monoxide releasing molecule 3
TUNEL: Terminal deoxynucleotidyl transferase dUTP nick end labeling
RAG: Regeneration associated genes
wnt4a: Wnt Family Member 4
stat3: Signal transducer and activator of transcription 3
dkk1b: Dickkopf Wnt Signalling Pathway Inhibitor 1
gsk3b: Glycogen Synthase Kinase 3 Beta
Tcf7l2: Transcription factor 7 like 2
Vasp: Vasodialator stimulated phosphoprotein
bZIP: Basic leucine zipper
Nrf2: Nuclear factor erythroid-derived 2-like 2
nfe2l2a: nfe2 like bZIP transcription factor 2a
SOD: Superoxide dismutase
GSH: reduced glutathione
GST: Glutathione S-Transferase
ARE: Antioxidant Response Element
WRE: Wnt Responsive Element
ER: Endoplasmic Reticulum
HGF: Hepatocyte Growth Factor
TSS: Transcription start site
ChIP: Chromatin Immunoprecipitation

## Declarations

### Ethics approval and consent to participate

The study was approved by the Institutional animal ethics committee (No. IAEC-04/BIOCHEM/CU/02-2022/03-SH) of Department of Biochemistry, University of Calcutta (Calcutta, India).

### Consent for publication

All authors gave their consent for publication of these research findings.

### Data availability

All data generated or analysed during this study are available from the corresponding author upon reasonable request.

### Authors’ contributions

RDS and SPH conceived and designed the study. RDS performed all experiments, acquired and analysed the data, and prepared the figures. AGC contributed to the conceptualization of the study and provided scientific input. AS contributed to the experimental methodology. RDS and SPH interpreted the data and wrote the manuscript. SPH supervised all aspects of the project. All authors read and approved the final manuscript.

## Acknowledgements

The authors gratefully acknowledge the research support provided through grants awarded to SPH by ANRF and DRDO, Govt. of India. RDS acknowledges the Council of Scientific and Industrial Research (CSIR), Government of India, for the Junior Research Fellowship (09/028(1146)-2020-EMR-I).

