## Supplementary Material for "Hmox1/CO regulates retina regeneration via modulation of Wnt/β-catenin signalling and induction of *vaspb* in zebrafish"

**Affiliations:**

**Email addresses of all authors**:

**ORCID IDs**:

Rinchen Doma Sherpa: https://orcid.org/0000-0002-9223-0289

Subhra Prakash Hui: https://orcid.org/0000-0001-5018-4918

Alok Ghosh Chaudhuri: https://orcid.org/0000-0003-4601-4486

**Supplementary Figure Legends**

**Supplementary Figure S.1:** qRT-PCR analysis of *hmox1b* in DMSO and ZnPPIX treated retina at 2dpi and 4 dpi (Mean ± SEM, n=5); $*p < 0.05$; $**p < 0.01$; *ns*, Non-significant; Mann-Whitney U test.

**Supplementary figure S.2: (**A-B) Immunohistochemical staining and quantification (B) of Hmox1$^+$ cells in Control morpholino (A.1) and Hmox1a morpholino (A.2) injected retina at 4 dpi; White arrowheads mark Hmox1$^+$ cells (Mean ± SEM, n=5); Magnification 100X; Scale bar 20 µm; $*p < 0.05$; $**p < 0.01$; *ns*, Non-significant; Mann-Whitney U test.

**Supplementary Figure S.3: (**A-B) Immunohistochemical staining and quantification (B) of PCNA$^+$ cells in Control morpholino (A.1) and Hmox1a morpholino (A.2) injected retina at 4dpi (Mean ± SEM, n=5); * represent injury point; Magnification 40X; Scale bar 50 µm. (C-D) Immunohistochemical staining and quantification (D) of average number of mislocalized HuC/D$^+$ cells at the injury epicenter in the INL and GCL in Control morpholino (C.1) and Hmox1a morpholino (C.2) injected retina at 28 dpi (Mean ± SEM, n=5); * represents injury point; White arrow head points towards migrating cells at the injury site, Magnification 40X; Scale bar 50 µm. $*p < 0.05$, $**p < 0.01$; *ns*, Non-significant; Mann-Whitney U test (B, D); INL = inner Nuclear Layer, GCL= ganglion cell layer.

**Supplementary Figure S.4:** (A-B) Immunohistochemical staining and quantification of EdU$^+$ cells in DMSO (A.1) and ZnPPIX (A.2) treated retina at 8 dpi (Mean ± SEM, n=5); * represents injury point; Magnification 40X; Scale bar 50 µm; *ns*, Non-significant; Mann-Whitney U test.

**Supplementary Figure S.5:** (A-B) Immunohistochemical staining and quantification of PCNA$^+$ cells in ZnPPIX+iCORM-3 (A.1), ZnPPIX+2 μM CORM-3 (A.2), ZnPPIX+ 5 μM CORM-3(A.3) and ZnPPIX+10 μM CORM-3(A.4) injected retina at 4dpi (Mean ± SEM, n=5); *represents injury point; Magnification 40X; Scale bar 50 μm. *$p < 0.05$; **$p < 0.01$; *ns*, Non-significant; Kruskal–Wallis with Dunn's multiple comparison test.

**Supplementary Figure S.6:** (A-B) Immunohistochemical staining and quantification of TUNEL$^+$ cells in DMSO (A.1), ZnPPIX (A.2) and ZnPPIX+CORM-3 (A.3) treated retinas at 1 dpi (Mean ± SEM, n=5); * represents injury point; Magnification 40X; Scale bar 50 μm; *ns*, Non-significant; One-way ANOVA with Tukey's multiple comparison.

**Supplementary Figure S.7:** (A-B) qPCR-based analysis of inflammatory genes at 1dpi (A) and anti-inflammatory genes at 2 dpi (B) in ZnPPIX and ZnPPIX+CORM-3 treated retinas (Mean ± SEM, n=5); *$p < 0.05$; **$p < 0.01$; *ns*, Non-significant; Mann-Whitney U test.

**Supplementary Figure S.8:** (A-B) Immunohistochemical staining and quantification (B) of β-catenin$^+$ cells in Control Morpholino (A.1) and Hmox1a morpholino (A.2) injected retinas at 4 dpi; * represents injury point; White arrowheads mark β-catenin$^+$ accumulation at the INL (Mean ± SEM, n=5); Magnification 40X; Scale bar 50 μm. *$p < 0.05$; **$p < 0.01$; *ns*, Non-significant; Mann-Whitney U test.

**Supplementary Figure S.9:** (A) qRT-PCR analysis of *nfe2l2a* & *nfe2l2b and nfe2l3* mRNA, time-course expression in regenerating retina (Mean ± SEM, n=5);

*$p < 0.05$; **$p < 0.01$; *ns*, Non-significant; One-way ANOVA with Dunnett's multiple comparison test.

**Supplementary figure S.10:** (A-C) Time course analysis of SOD (A) and GSH (B) activities and GST (C) levels at different time points following retinal injury (Mean ±

SEM, n=3) (D) Multiple sequence alignment of NRF2-bZIP DNA binding domain between zebrafish (*Danio rerio*), mouse (*Mus musculus*), and human (*Homo sapiens*). **p* < 0.05; ***p* < 0.01; ****p*<0.001; *ns*, Non-significant; One-way ANOVA with Dunnett's multiple comparison test.

**Supplementary Table S.1**: *In silico* predictions of Tcf/Lef binding motifs (Human/Mouse TCF7L2/Tcf7; JASPAR) on *vaspb* upstream regulatory regions.

**Supplementary Table S.2**: *In silico* predictions of Nrf2 binding Antioxidant Response Elements (AREs) in the upstream regulatory regions of zebrafish regeneration associated genes

**Supplementary Figure S.1**

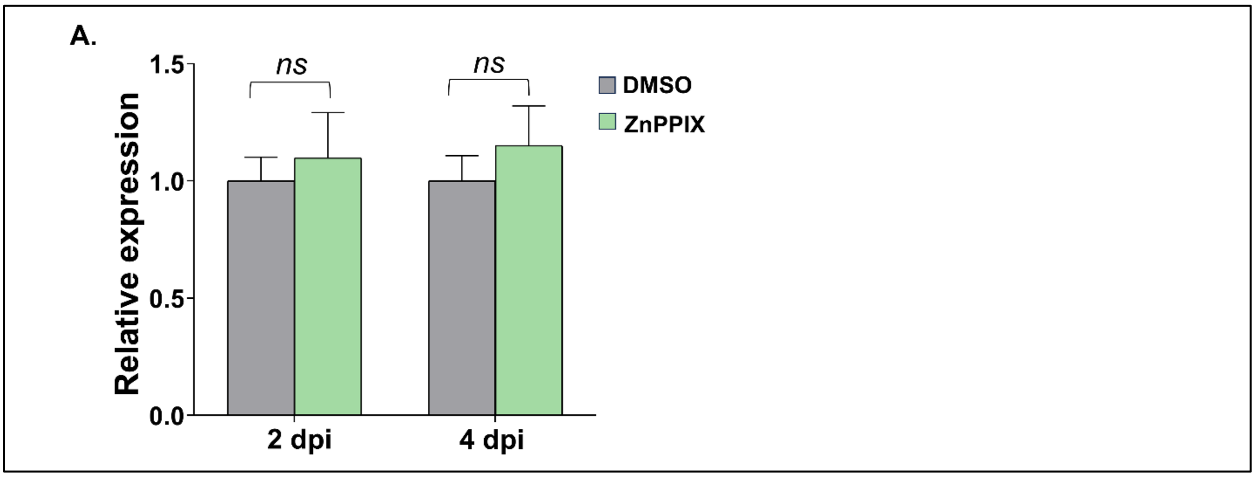


**Supplementary Figure S.2**

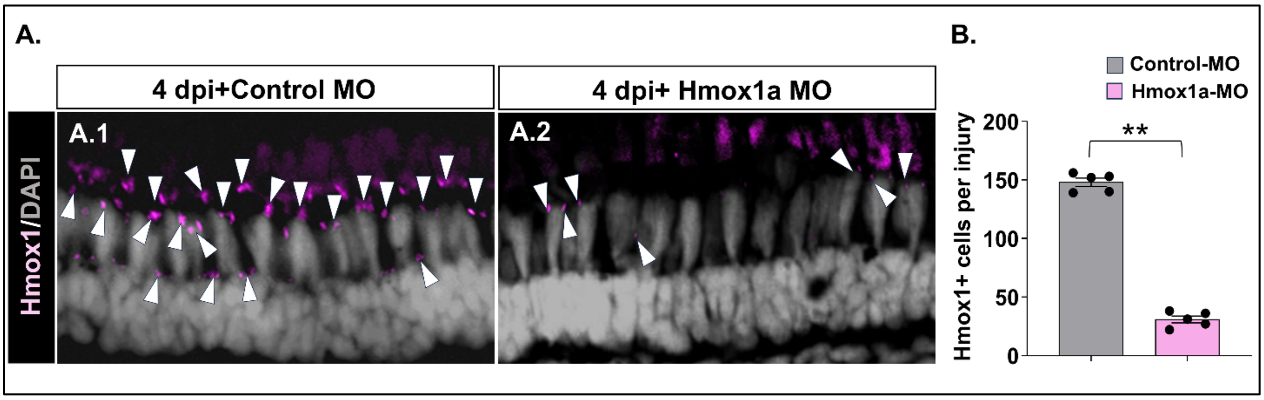

### Supplementary Figure S.3

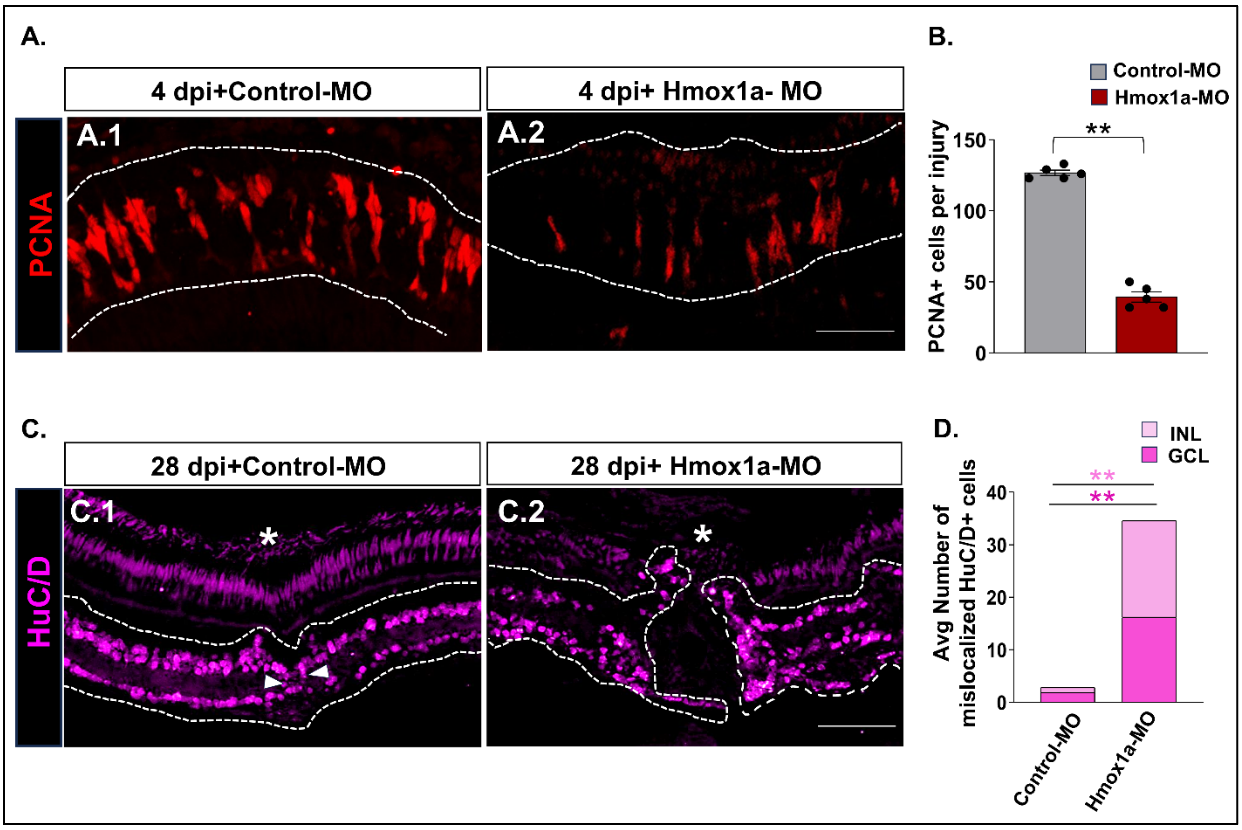


### Supplementary Figure S.4

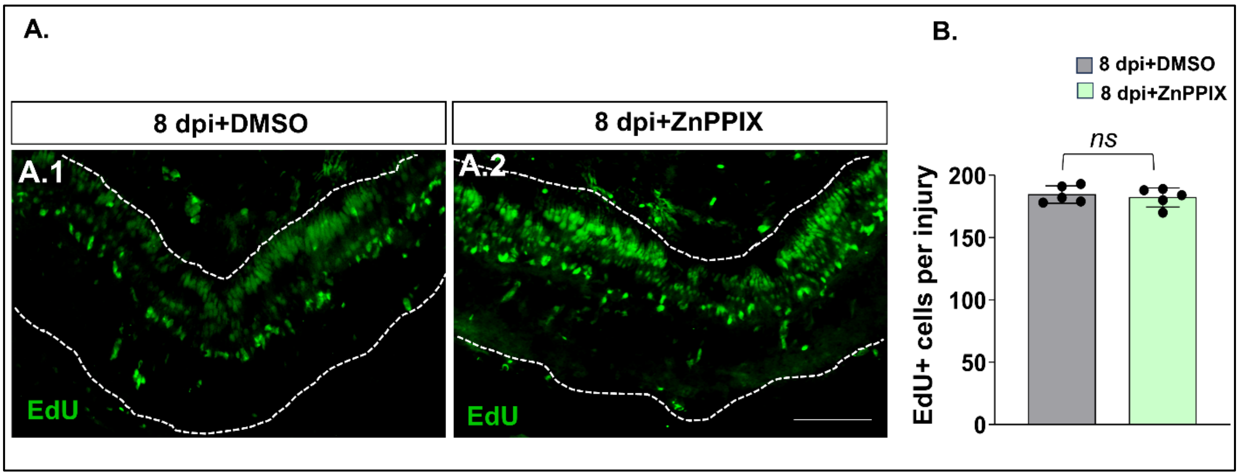

### Supplementary Figure S.5

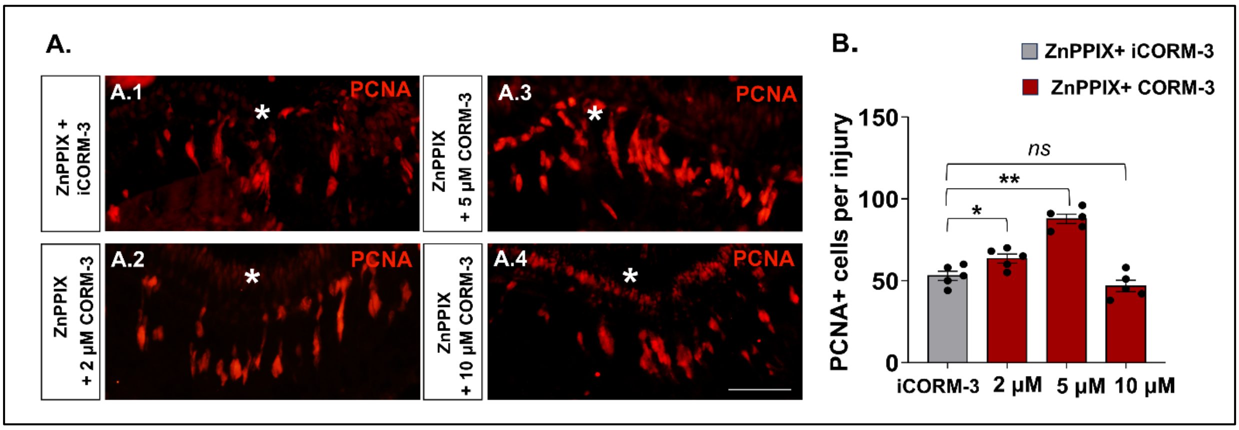


### Supplementary Figure S.6

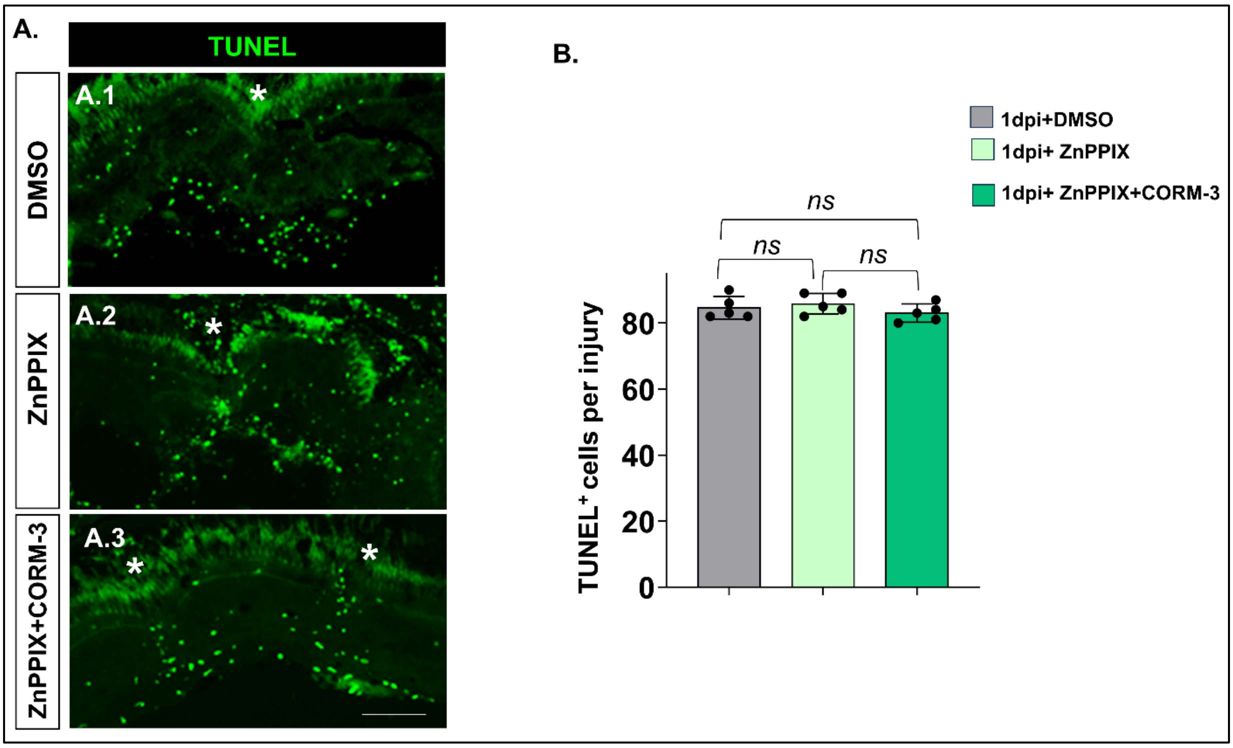

### Supplementary Figure S.7

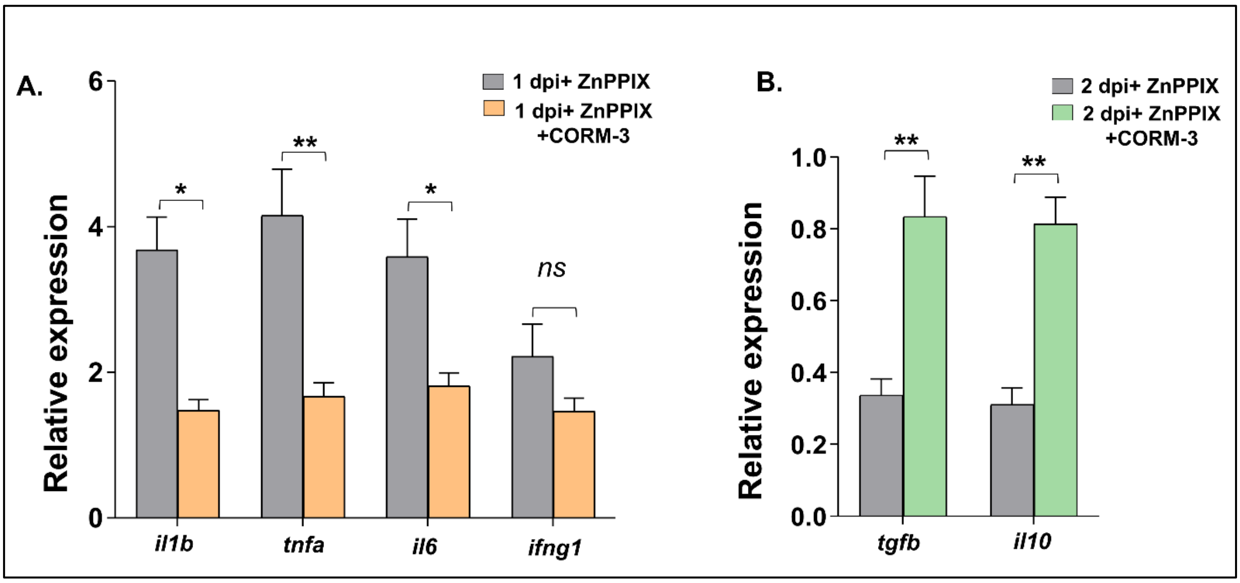


### Supplementary Figure S.8

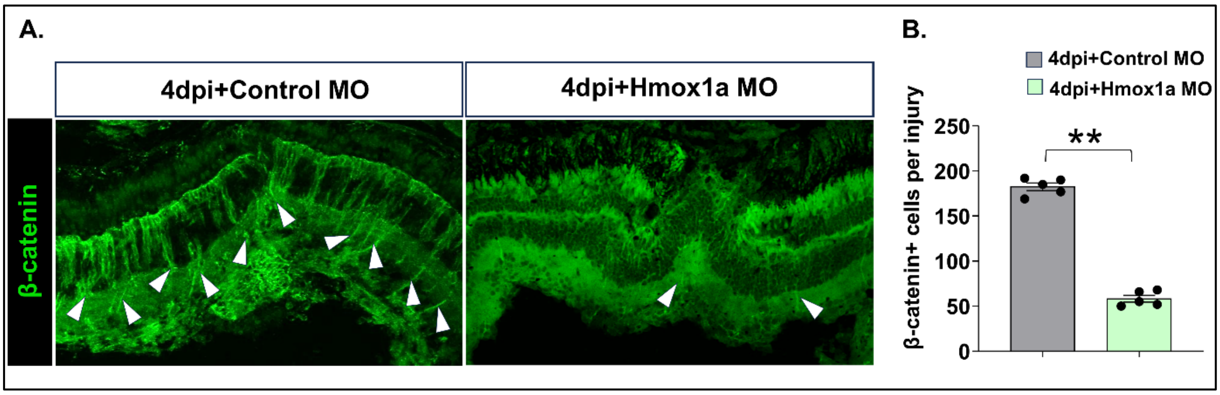

### Supplementary Figure S.9

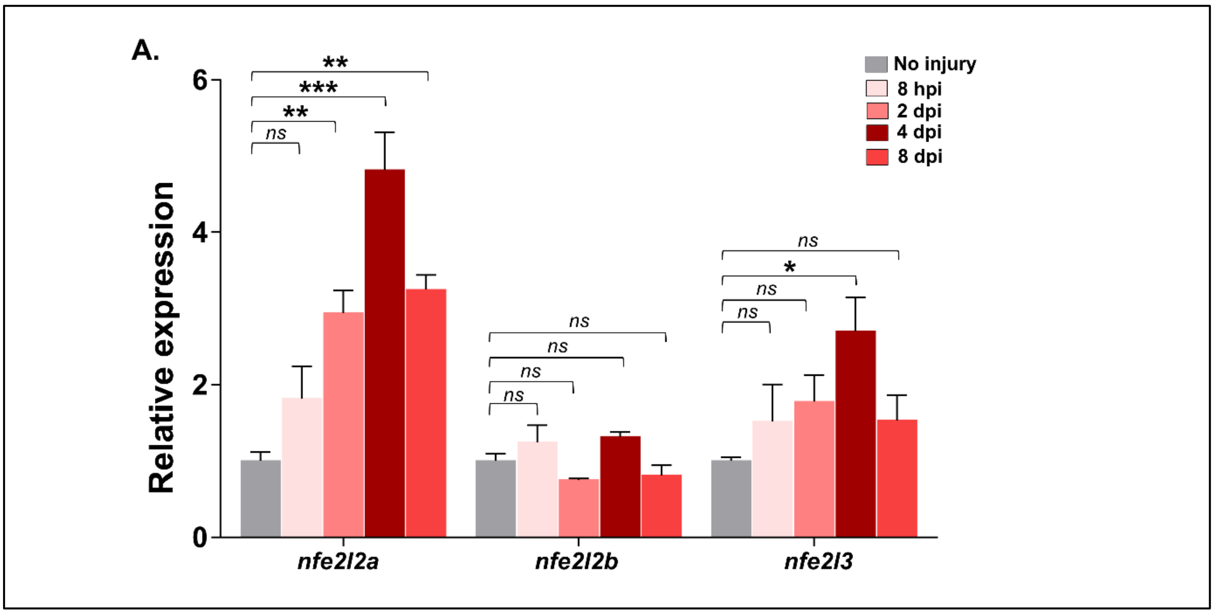


### Supplementary Figure S.10

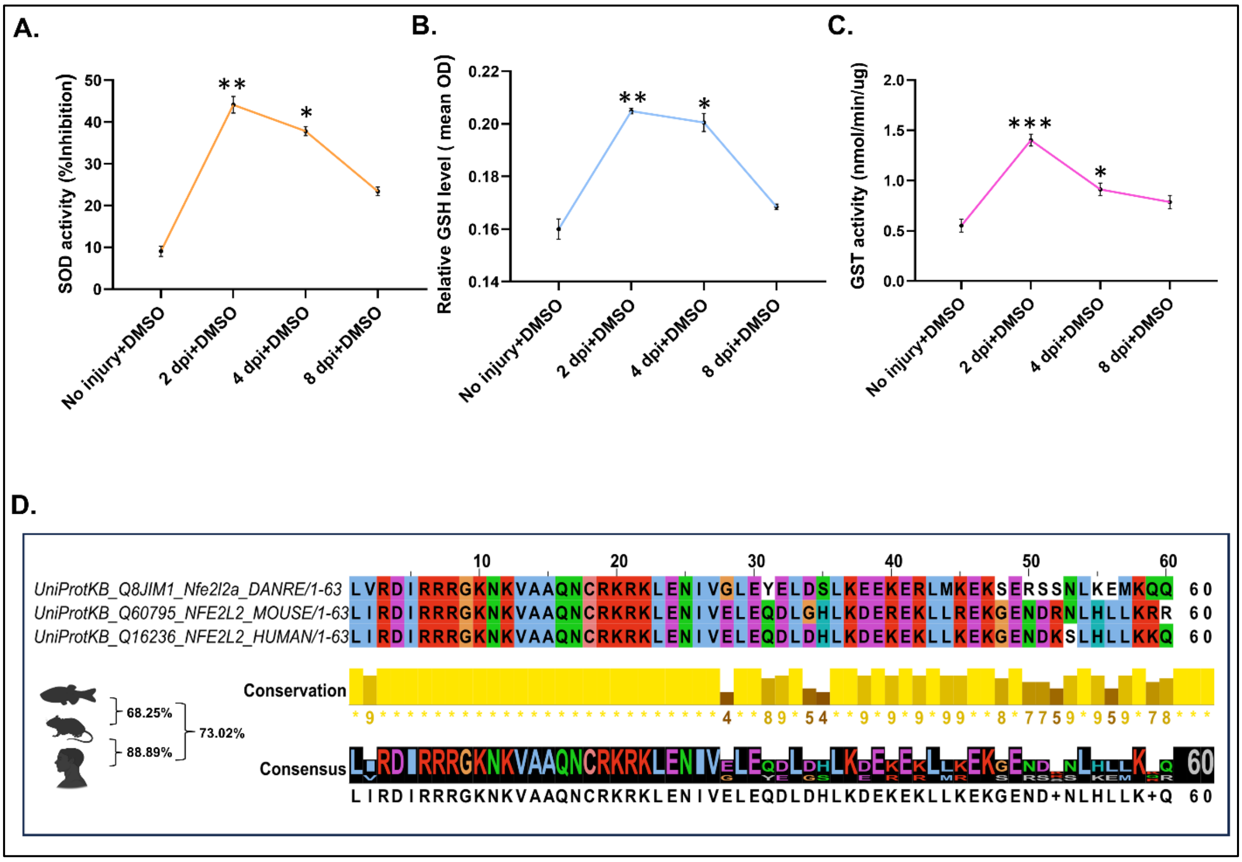

### Supplementary Table S.1

FIMO identified Tcf7/TCF7L binding regions on zebrafish *vaspb* (p<0.0001, q<0.32)

| JASPAR motif_id | Species | motif_alt_id | sequence_name | start | stop | strand | score | p-value | q-value | matched_sequence |
|---|---|---|---|---|---|---|---|---|---|---|
| **MA0523.1** | ***Homo sapien*** | **TCF7L2** | 18 | 1463 | 1476 | + | 13.0182 | 2.19E-05 | 0.304 | CAACATGAAAGCAC |
| **MA0523.1** | ***Homo sapien*** | **TCF7L2** | 18 | 3206 | 3219 | + | 12.6 | 2.92E-05 | 0.304 | AGACTTCAAAGTAG |
| **MA0523.1** | ***Homo sapien*** | **TCF7L2** | 18 | 9385 | 9398 | - | 11.3273 | 6.01E-05 | 0.304 | AATGATCAAAAGGA |
| **MA0523.1** | ***Homo sapien*** | **TCF7L2** | 18 | 7529 | 7542 | + | 10.9273 | 7.43E-05 | 0.304 | GGAATTCAAAGGGC |
| **MA0523.1** | ***Homo sapien*** | **TCF7L2** | 18 | 4692 | 4705 | + | 10.7636 | 8.13E-05 | 0.304 | AAATATGAAAGGGT |
| **MA0523.1** | ***Homo sapien*** | **TCF7L2** | 18 | 4027 | 4040 | - | 10.7273 | 8.28E-05 | 0.304 | GAAGTTCAAAAACG |
| **MA0769.1** | ***Mus Musculus*** | **Tcf7** | 18 | 1463 | 1474 | + | 9.5 | 6.04E-05 | 0.319 | CAACATGAAAGC |
| **MA0769.1** | ***Mus Musculus*** | **Tcf7** | 18 | 9322 | 9333 | - | 12.1613 | 1.80E-05 | 0.319 | ATAGATTAAAGG |
| **MA0769.1** | ***Mus Musculus*** | **Tcf7** | 18 | 9387 | 9398 | - | 10.9839 | 3.28E-05 | 0.319 | AATGATCAAAAG |
| **MA0769.1** | ***Mus Musculus*** | **Tcf7** | 18 | 3206 | 3217 | + | 9.80645 | 5.33E-05 | 0.319 | AGACTTCAAAGT |

### Supplementary Table S.2

FIMO identified Antioxidant Response Elements on Wnt-Responsive genes (p<0.0001, q<0.25)

| Gene | motif_id | motif_alt_id | chromosome number | start | stop | strand | score | p-value | q-value | matched_sequence |
|---|---|---|---|---|---|---|---|---|---|---|
| ***dkk1b*** | **MA0150.1** | **NFE2L2** | 12 | 9606 | 9616 | + | 14.6122 | 7.19E-06 | 0.144 | ATGATTCAGCA |
| ***dkk1b*** | **MA0150.1** | **NFE2L2** | 12 | 9272 | 9282 | - | 12.4898 | 2.91E-05 | 0.25 | CTGACATAGCA |
| ***dkk1b*** | **MA0150.1** | **NFE2L2** | 12 | 3916 | 3926 | - | 11.9388 | 3.76E-05 | 0.25 | ATGACAAAGCT |
| ***gsk3bb*** | **MA0150.1** | **NFE2L2** | 1 | 877 | 887 | + | 14.5204 | 7.61E-06 | 0.152 | GTGACACGGCA |
| ***wnt4*** | **MA0150.1** | **NFE2L2** | 11 | 6868 | 6878 | + | 14.8163 | 5.85E-06 | 0.117 | GTGACTGAGCA |

**Key Resource Table**

| Reagent/ Resource | Source | Identifier |
|---|---|---|
| **Antibodies** | | |
| Rabbit polyclonal anti HO-1 | Enzo Biochemc | Cat# BML-HC3001-0100 |
| Mouse monoclonal anti-PCNA | Abcam | Cat# ab29 (RRID:AB_303394) |
| Mouse monoclonal anti-HuC/D | Invitrogen | Cat# 16A11 |
| Rabbit polyclonal anti-βcatenin | Santa Cruz | Cat# sc-7199 |
| Rabbit polyclonal VASP | Proteintech | Cat# 13472-1-AP |
| Goat anti-rabbit Alexa Fluor 488, IgG (H+L) | Jackson Immunoresearch | Cat# 111-545-003 (RRID:AB_2338046) |
| Goat anti-mouse Alexa Fluor 488, IgG (H+L) | Jackson Immunoresearch | Cat# 115-545-062 (RRID:AB_2338845) |
| Goat anti-rabbit Alexa Fluor 594, IgG (H+L) | Jackson Immunoresearch | Cat#111-584-144 |
| Goat anti-mouse Alexa Fluor 594, IgG (H+L) | Jackson Immunoresearch | Cat# 115-585-566 |
| Rabbit polyclonal IgG | Abcam | Cat# ab37415 |
| **Reagents** | | |
| ZnPPIX | Cayman Chemical | Cat# 15442-64-5 |
| CORM-3 | Merck | Cat# SML0496 |
| Tissue Freezing Medium | Leica Biosystems | Cat# 14020108926 |
| DAPI (4’,6- diamidino-2-phenylindole dihydrochloride) | Sigma-Aldrich | Cat# D9542 |
| TRIzol Reagent | Thermo Fisher Scientific | Cat# 15596026 |
| Dimethyl Sulfoxide (DMSO) | Sigma-Aldrich | Cat# D4540 |

| | | |
|---|---|---|
| Fetal Bovine Serum | Thermo Fisher Scientific | Cat# 10082139 |
| Dabco 33-LV (1,4-Diazabicyclo[2.2.2]octane solution) | Sigma | Cat# 290734 |
| **Critical Commercial Assays** | | |
| GoScript™ Reverse Transcriptase Kit | Promega | Cat# A5003 |
| GoTaq Green Master Mix | Promega | Cat# M7122 |
| PrimeSTAR GXL DNA Polymerase | Takara Clonetech | Cat# R051A |
| SYBR Select Master Mix | Thermo Fisher Scientific | Cat# 4472908 |
| *In Situ* Cell Death Detection Kit, Fluorescein | Roche | Cat# 11684795910 |
| Click-iT™ EdU Cell Proliferation Kit for Imaging, Alexa Fluor™ 488 dye | Invitrogen | Cat# C10337 |
| Pierce™ Bradford Plus Protein Assay Reagent | Thermo Scientific | Cat# 23238 |
| Pyrogallol | SRL | Cat# 71646 |
| 1 chloro-2,4- Dinitrobenzene | SRL | Cat# 42687 |
| 5,5'-Dithiobis (2-Nitro Benzoic Acid) extrapure (DTNB), 98% (Ellmans Reagent) | SRL | Cat# 69-78-3 |
| Glutathione Reduced (GSH) | SRL | Cat# 48938 |
| Trichloroacetic Acid 10% solution | SRL | Cat# 31445 |
| EpiQuik™ Tissue Chromatin Immunoprecipitation (ChIP) Kit | Epigentek | Cat# P-2003-2 |
| **Oligonucleotides** | | |
| *hmox1a* morpholino: 5′-TGGAGTCCATCTTTGTGCTGTAGA -3′ | Gene Tools | N/A |

| | | |
|---|---|---|
| Primers for qRT-PCR, see Oligonucleotide Table | | N/A |
| **Software** | | |
| Fiji (ImageJ) (RRID:SCR_002285) | | https://fiji.sc/ |
| Primer3 (RRID:SCR_003139) | | https://primer3.ut.ee/ |
| FV31S-SW software | | N/A |

#### Oligonucleotide Table

| Gene | Forward primer 5'→3' | Reverse primer 5'→3' | Experiments |
|---|---|---|---|
| ***hmox1a*** | TCTACAGCACAAAGATGGACTCCA | ACAGCAGAAGCTTGTACTGTGT | qRT_PCR |
| ***hmox1b*** | GCAGTGATCTGTCTGAACAG | GCTTGTACTGTGTTTGTGTG | qRT_PCR |
| ***hmox2a*** | TGCGAAATCAGAGAAGAGCCTGA | AGTGGTGCCAAGCTTAAAAAGTTC | qRT_PCR |
| ***hmox2b*** | CAAACAGCGGTTGTTGTGTTT | CAAACAGCGGTTGTTGTGTTT | qRT_PCR |
| ***blvra*** | TGTGCTTACAGAGTTTGCCTC | AGTTAAGCTGCCCTGTCAAGA | qRT_PCR |
| ***blvrb*** | AGAACTTGCTACGTCTGTCTCTT | ACCAGCAATATGTGGAGGCAT | qRT_PCR |
| ***il1b*** | TGAAGCACATCAAACCCCAATC | TAAGACGGCACTGAATCCACCA | qRT_PCR |
| ***tnfα*** | TTGGGATCATTTTGGCTGTGG | TCCATCTTGTTGATTGCCCTG | qRT_PCR |
| ***il6*** | TCTGTACAAGGACGTGAAGACAC | TGATCCTGACCCCTTCAAATG | qRT_PCR |
| ***ifnγ1.2*** | TGAAGCTACAAAGGAGAGGCT | AACAAAGCCTTTCGCTGGAC | qRT_PCR |
| ***il10*** | AAGCGGGATATGGTGAAATG | AGACCCCCTTTTCCTTCATC | qRT_PCR |
| ***tgfβ*** | TACTGCGACTGTAAAGCAAACCA | TCACCATCATATCTGCCAGACC | qRT_PCR |
| ***ascl1a*** | ACTCCGCGAAATGGACATCA | TGCTTTGACGCGGACTTGTT | qRT_PCR |
| ***pouf53*** | TATTACGAGCAGAGTCCACCAC | AGCAAGTTAGCTGGTGAGATGA | qRT_PCR |
| ***sox2*** | ATGATCAGTATGTACCTGCCCG | ATCGTGCCGTTAATCGTCGT | qRT_PCR |
| ***stat3*** | CTGTAGGAACATGGGATCAGGT | TCTGGCAACCGGAGTAGTTTAC | qRT_PCR |
| ***lin28*** | ATAAAGATGCCCCCGGCAAATC | TGCTCTGATGGACGAAGACATCCA | qRT_PCR |
| ***wnt4a*** | CAATGCGAGCAACTGGCTATAC | AATGCAGCTTCCCTCGTACCTT | qRT_PCR |

| | | | |
|---|---|---|---|
| ***wnt8b*** | TTGACCATTCATTGGGTTTTGAC | AACATGTTGCATTTCTCAAGGC | qRT_PCR |
| ***pdgfaa*** | CCAGACGCAAGAGAAGCATT | GATCACCGTACGGGTCTTACA | qRT_PCR |
| ***fgf*** | TTCCTTGCCATGAACGCAGA | TAGTTGTTGGACTCCAGGCG | qRT_PCR |
| ***notch1a*** | AGGCGGAGTTATGAACCGTT | ACACCTGCATGATGCTTCTC | qRT_PCR |
| ***mmp9*** | TTTGCGACTCAAAATGAGACTTGG | AACGCTTCAGATACTCATCCG | qRT_PCR |
| ***shha*** | AAGCCCACATTCATTGCTCT | CCTTCTGTCCTCCGTCCTG | qRT_PCR |
| ***vaspa*** | TGCCTTTCAGTTCCCACTGT | AGCTACATGCACGAAAACAGA | qRT_PCR |
| ***vaspb*** | ATCTAGTCCACTTGCCAGGAT | ACTCCTGAATAAAGGCTCCGA | qRT_PCR |
| ***nfe2l2a*** | CACACCTTAAAATAGCAGTTGTC | AAGCACCACTGGAAAATTTAGA | qRT_PCR |
| ***nfe2l2b*** | TGCATGGAAGAGAAAGAGAGGAA | GGATTGTACGAGTTCCCATGCTC | qRT_PCR |
| ***nfe2l3*** | ACAGAATTGTCGCAAGAGGAAGC | CATAGTCCGCGAGAGTGCTTC | qRT_PCR |
| ***vaspb promoter*** | GAGGAATGCAGAAGAGCATCTCTAACC | AGATGGGCTCCAGCGATTGCCA | CHIP_PCR |
| ***actb2*** | ATCAAGATCATTGCTCCCCCT | GGTTGGTCGTTCGTTTGAATCT | qRT_PCR |
